# Interleukin-17 signaling influences CD8^+^ T cell immunity and tumor progression according to the IL-17 receptor subunit expression pattern in cancer cells

**DOI:** 10.1101/2022.12.16.520754

**Authors:** Constanza Rodriguez, Cintia L. Araujo Furlan, Jimena Tosello Boari, Sabrina N. Bossio, Santiago Boccardo, Laura Fozzatti, Fernando P. Canale, Cristian G. Beccaria, Nicolás G. Nuñez, Danilo G. Ceschin, Eliane Piaggio, Adriana Gruppi, Carolina L. Montes, Eva V. Acosta Rodríguez

## Abstract

The role of IL-17 mediated immune responses in cancer is conflicting as pre-clinical and clinical results show tumor-promoting as wel as tumor-repressing functions. Herein, we used syngeneic tumor models from different tissue origins as a tool to evaluate the role of IL-17 signaling in cancer progression, dissecting the effects in cancer cell growth and tumor immunity. We show that absence of IL-17RA or IL-17A/F expression in the host has contrasting effects in the *in vivo* growth of different tumor types. We observed that lack of IL-17A/F-IL-17RA signaling in host cells changed the expression pattern of several mediators within the tumor microenvironment in a cancer-type specific manner. Deficiencies in host IL-17RA or IL-17A/F expression resulted in reduced antitumor CD8+ T cell immunity in all cancer models and in tumor-specific changes in several lymphoid cell populations. These findings were associated to particular patterns of expression of cytokines (IL-17A and IL-17F) and receptor subunits (IL-17RA, IL-17RC and IL-17RD) of the IL-17 family in the injected tumor cell lines that, in turn, dictated tumor cell responsiveness to IL-17. We identified IL-17RC as an important determinant of the IL-17-mediated transcriptional response in tumor cells and; consequently, as a predictive biomarker of the overall effect of IL-17 signaling in tumor progression. Our findings contribute to unraveling the molecular mechanisms underlying the divergent activities of IL-17 in cancer and provide rational targets for immunotherapies based on personalized approaches.

## INTRODUCTION

The interleukin 17 (IL-17) family comprises six cytokines named IL-17A (or IL-17), IL-17B, IL-17C, IL-17D, IL-17E (also called IL-25) and IL-17F. Among them, IL-17A and IL-17F stand out as they are coordinately produced by many immune cells and promote inflammation through the induction of cytokines, chemokines, growth factors, metalloproteinases (MMP) and antimicrobial peptides, among others (1). Accordingly, IL-17A and IL-17F are associated to the resolution of infections with a vast array of microorganisms as well as involved in the promotion of autoimmune diseases (2). In the context of malignancy, IL-17A has been detected in serum and tumor-associated fluids while IL-17-producing cells have been found in blood, lymphoid organs and tumor tissues from various cancer types, both in experimental models and humans (3, 4). Despite intensive research in the last years, the function of IL-17-mediated responses in cancer settings remains conflicting as pre-clinical and clinical results show tumor-promoting as well as tumor-repressing effects (5). Responsiveness to IL-17 is mediated by the expression of surface receptors formed by homo- or hetero-complexes of two or three subunits of the IL-17 receptor (IL-17R) family (2). The IL-17R family is composed of 5 members, named from IL-17RA to IL-17RE. IL-17RA was first discovered as the receptor for IL-17A, IL-17F and the heterodimer IL-17A/F. Later, IL-17RA was reported to also bind to IL-17E and IL-17C; so, it is now considered as a common subunit of the IL-17-receptor family. The complex formed by IL-17RA and IL-17RC is considered the canonical receptor that mediates the signaling by IL-17A and IL-17F (6). However, IL-17RA can also complex with IL-17RD to allow IL-17A, but not IL-17F, signaling (7, 8). Of note, transcriptome analyses demonstrated that, when compared with the canonical IL-17RA/RC receptor complex, the IL-17RA/RD complex triggers overlapping as well as unique gene regulation after IL-17A ligation (9). Furthermore, a recent report demonstrated that IL-17RC is able to form homodimeric complexes that participate in IL-17RA-independent IL-17 signaling (10). Thus, the combination of IL-17R subunit expression together with different cytokine/receptor affinities may dictate the responsiveness to the IL-17 cytokine family. In cancer, this complexity of the IL-17A/IL-17RA signaling may explain the divergences in the effect of this pathway in each context.

A systemic analysis of a large set of published results concluded that IL-17A was associated with poor outcome in human cancers while Th17 cells were associated with better prognosis, although tumor-type particularities were observed (11). IL-17 tumor-promoting functions include direct effects in tumor cells, and indirect effects through the modulation of stromal cells and the immune cell infiltrate of the tumor microenvironment (12). In fact, IL-17A signaling directly promotes self-renewal, survival and therapy resistance in cancer cells of many histological origins (13–15). Furthermore, IL-17A induces the expression of MMPs and pro-angiogenic factors such as VEGF in tumor cells and cells from the tumor microenvironment, promoting angiogenesis, cell invasion and metastasis (16, 17). Finally, IL-17A has been shown to reshape the tumor immune infiltrates by sustaining the recruitment of myeloid cells that support immunosuppression and, consequently, tumor progression (18–20). Given these antecedents, many researchers have postulated the IL-17/L-17R axis as a potential novel immunotherapeutic target for cancer (3, 12, 21, 22). Notwithstanding the probable involvement of IL-17A in tumorigenesis, diverse studies have showed that IL-17A and IL-17-producing cells may potentiate anti-tumor immunity through the recruitment and activation of effector immune cell subsets (4). In this regard, IL-17 mediated dendritic cell and neutrophil recruitment and activation leading to enhanced anti-tumor CD8+ T cell immunity in certain cancer settings (23–25). Furthermore, IL-17A promoted CXCL9 and CXCL10 production and favored the recruitment of effector T cells and NK cells, sustaining Th1-type responses (26–28). In addition, transfection of IL-17A into a Meth-A fibrosarcoma favored tumor cell elimination by the induction of class I and II MHC molecules (29). In all, these findings highlight the dichotomous nature of the IL-17/IL-17R pathway in malignancy. In fact, accumulating evidences support the hypothesis that IL-17A shows pro-tumor properties at the early stages of carcinogenesis, whereas its role in established tumors is variable depending on the tumor-specific context (5). As the malignant clones are the most particular components of the tumor microenvironment, whether tumor cells are able (or not) to respond to IL-17 itself may be a key determinant for the tumor-promoting or suppressing effects of this cytokine.

Herein, we used syngeneic tumor models from different tissue origins as a tool to evaluate the role of IL-17/IL-17F signaling in cancer progression, dissecting the effects in cancer cell growth and in tumor immunity. We determined that absence of IL-17RA or IL-17A/F expression in the host had contrasting effects in the *in vivo* growth of different tumor types. We observed that lack of IL-17A/F-IL-17RA signaling in host cells changed the expression pattern of several mediators within the tumor microenvironment in a cancer-type specific manner. Remarkably, deficiencies in host IL-17RA or IL-17A/F expression had a particular impact in the antitumor CD8+ T cell response, as well as in the recruitment of different lymphoid, rather than myeloid cell populations depending on the tumor model. Interestingly, these *in vivo* findings associated to particular patterns of expression of cytokines (IL-17A and IL-17F) and receptor subunits (IL-17RA, IL-17RC and IL-17RD) of the IL-17 family present in the injected cell lines that, in turn, dictated tumor cell responsiveness to IL-17. We identified IL-17RC as an important determinant of the IL-17-mediated transcriptional response in tumor cells, and consequently, as a predictive biomarker of the overall effect of IL-17 signaling in tumor progression. Our findings have relevance in unraveling the molecular mechanisms underlying the divergent activities of IL-17 in cancer and provide rational targets for immunotherapies based on personalized approaches.

## MATERIALS AND METHODS

### Cell lines

Melanoma (B16F10.SIY.GFP), Fibrosarcoma (MC57.SIY.GFP), Acute Myeloid Leukemia (C1498.SIY.GFP) and Lymphoma (EL4.SIY.GFP) tumor cells were engineered to express GFP fused in frame with the model antigen SIYRYYGL, which is recognized by CD8+ T cells in the context of K^b^ (30–32). Thus, all these tumor models enable monitoring of SIY antigen-specific T cell responses in tumor-bearing hosts. Cells were maintained in Complete Medium: DMEM (Cornell) supplemented with 10% (v/v) heat-inactivated fetal bovine serum (FBS) (GIBCO or Natocor), 1 mM L-glutamine (GIBCO), 25 mM Hepes (Gibco) and 55 μM β-mercaptoethanol (GIBCO) and a mixture of streptomycin (100ug/mL) and penicillin (100 units/mL) (GIBCO). To minimize sources of cell variability, our protocol included the periodically Mycoplasma testing as well as the selective purification of GFP+ SIY-expressing cells to achieve a homogeneous cell population.

For IL-17A and IL-17F stimulation experiments, B16F10.SIY.GFP, MC57.SIY.GFP, C1498.SIY.GFP and EL4.SIY.GFP cells were cultured in serum reduced (1%FBS) complete medium, and stimulated with mouse recombinant IL-17A (200ng/ml) (Shenandoah Bt, Catalog: 200-59-100ug), or IL-17F (Shenandoah Bt, Catalog: 200-62-100ug). Each condition was assessed in triplicates.

For IL-17RC blocking assay, B16F10SIY.GFP and EL4.SIY.GFP cells were plated in serum reduced complete medium (1% FBS, DMEM) and stimulated for 24 hours with IL-17A (200ng/mL, cat: 200-59-100ug, Shenandoah) and anti-IL-17RC (5ug/mL, cat: PA5-47286, Invitrogen). Goat IgG (5ug/mL, cat: BAF108, R&D) was used as a control. The following day, cells were harvested for RNA extraction and culture supernatants were collected.

### Mice

C57BL/6 (B6) WT mice were inbred at our Institute. IL-17RAKO mice were provided by Amgen Inc. (Master Agreement N° 200716544-002), while IL-17A and IL-17F double knock out (IL-17A/F DKO) mice were originally provided by Dr Immo Prinz (33). Age and sex matched female or male mice from 6 to 12 weeks of age were used for the experiments carried out in this work. Genetically modified animals were regularly checked by PCR assays for strain control. The institutional animal facility follows the recommendations of the Guide for the Care and Use of Experimental Animals, published by the Canadian Council for the Protection of Animals. The protocol was approved by the IACUC of Facultad de Ciencias Químicas, Universidad Nacional de Córdoba under number RD 732/18.

### Experimental mouse tumor models

For tumor establishment, cultured cells were trypsinized (if required), washed and counted in Neubauer chamber with trypan blue. 2×10^6^ B16F10.SIY.GFP, MC57.SIY.GFP and C1498.SIY.GFP or 0.5×10^6^ EL4.SIY.GFP cell suspensions were injected subcutaneously (s.c.) in 200uL of sterile PBS to the different mouse strains. Tumor volume, was determined by caliper 3 times per week using the formula V= (smallest diameter (d) x largest diameter (D)^2^) x 0.5 (Faustino-Rocha et al., 2013). Mice were sacrificed if they showed signs of marked weight loss, tumor ulceration or tumor volume greater than 2000mm^3^.

### Cell preparation and flow cytometry

Tumors were obtained from WT, IL-17RA KO or IL-17A/F DKO mice injected with B16F10-SIY.GFP and EL4-SIY.GFP and disaggregated first mechanically and then enzymatically with 2mL of RPMI containing 2 mg/mL collagenase IV (Roche, cat: 11088882001) and 50 U/mL DNase I (Roche, cat:4536282001). Tumor suspensions were filtered with 70 μm cell strainers (BD Biosciences), washed and kept in cold RPMI-2% FBS. The number of cells was determined by counting in a Neubauer chamber using Türk’s solution.

Tumor cell suspensions were treated with FcR Blocker (Stem Cell Technologies) for 10min at RT. Afterwards, cells were incubated during 20 min at 4°C with a combination of the required antibodies (Supplementary Table 1, 2 and 3). When staining involved the identification of SIY-specific CD8+ T lymphocytes, a previous incubation step with a SIY dextran (SIYRYYGL) conjugated with PE fluorochrome (Immudex) was added. A viability dye Fixable Viability Stain 700 (BD) or Fixable Aqua Dead Cell Stain Kit, for 405 nm excitation (Invitrogen), was included in the staining (Supplementary Table 3).

All samples were acquired on BD FACSCanto II, BD LSR Fortessa X20 (BD Biosciences) or Attune^Nxt^ (Life Technologies) flow cytometers. The data generated was analyzed using FlowJo software, in all cases, gating strategy included event selection according to size and granularity based on FSC-A vs SSC-A dot plot. Also death cells, doublets and aberrant events due to laser fluctuation over time were excluded. An additional step was added when evaluating tumor infiltration by eliminating GFP+ CD45-tumor cells (Supplementary figure 2). Unsupervised analysis was performed on live CD45+ cells using UMAP and FlowSOM Flowjo pluggins (34, 35).

### Angiogenesis Protein Array

To evaluate angiogenesis related proteins, tumor lysates were prepared according to manufacturer’s specifications using B16.SIY and EL4.SIY tumors obtained from WT, IL-17A/F DKO and IL-17RAKO hosts at 18-19 dpi. Protein content in each tumor lysate was quantified using Bradford protein assay. Final samples corresponding to each tumor line and mouse strain to be probed in the membrane were prepared by pooling equal amounts of proteins from tumor lysate of all replicates to obtain 4000ug of proteins per 1000uL of solution. The expression of angiogenesis-related proteins was detected by using the Proteome Profiler Mouse Angiogenesis Array Kit (ARY015; R&D Systems) following the manufacturer’s protocols. Developed films were scanned and analyzed by quantifying the mean spot pixel densities using the NIH ImageJ software. Corresponding signals on different arrays were normalized by using internal controls.

### Cell viability and proliferation assays

To determine proliferation by MTT assay (Kumar et al., 2018), cells were cultured in serum reduced conditions with IL-17A or IL-17F as previously described. After 24h of culture, MTT solution was added to each well at a final concentration of 0.5mg/ml. The plate was incubated for 4h and then revealed by adding DMSO. Proliferating cell numbers were obtained by reading absorbance at 570nm 15min later on plate reader SYNERGY HT (Biotek) instrument.

### Wound healing assay

Cell line monolayers were cultured in serum-reduced conditions with (or without) IL-17A (200ng/mL) and IL-17F (200ng/mL). Two wounds were performed with a tip per cell monolayer. Each cell line and culture condition were assayed in duplicates. For quantification, wound closure was assessed at 0, 6 and 12h post stimulus, taking serial photographs with a Leica DMI 8 microscope using the Mark and Find function. Photographs were analyzed with Fiji - ImageJ software using PHANTAST plugging. The percentage of closure was obtained by calculating the ratio between the free area (fa) at each time point evaluated and the free area at time zero (fa0), which was defined as 0% closure. The formula used was: 100-(fa*100/fa0) (Yue et al., 2010).

### Protein quantification

Quantifications in culture supernatants, plasma and tumor lysates were performed by flow cytometry using a multiplex bead-based system (LEGENDPlex Mouse Th Cytokine Panel and Mouse Inflammation Panel kits, Biolegend) that allowed the simultaneous determination of the concentration of several cytokines. Samples were assessed according to the manufacturer’s instructions and acquired on an Attune^NxT^ flow cytometer. The files generated were analyzed according to fluorescence intensity in FlowJo software. Alternatively, in the indicated cases, cytokines and chemokines were quantified by ELISA using commercial kits (Ready-SET-Go!, eBioscience or Prepotech) following the manufacturer’s specifications. In all cases, analysis of the calibration curves and extrapolation of the concentration was performed using GraphPad Prism software.

### Western blot Analysis

Whole cell extracts and nuclear extracts were obtained after stimulation with IL-17A (200 ng/mL) at the indicated times. Whole cell lysates were obtained using RIPA buffer. Nuclear extracts were obtained after treating cells with Buffer A (10 mM HEPES; 10 mM KCl; 1 mM EDTA; 1.5 mM MgCl; 5 mM NaF), discarding the cytoplasmic protein fraction, and then, Buffer B was added (20 mM Hepes; 5mM NaF; 0.4 M NaCl). Finally, samples were centrifuged at 13,000g for 5min and the nuclear protein fraction was collected from the supernatant. The extracts were stored at −80°C until processing. Protein samples were quantified using Bradford assay and 40 ug were treated with sample buffer (0.125M Tris-Hcl pH 6.8, SDS 2.5 %, 25 % β-mercaptoethanol, 25 % Glycerol, 0.1 mg/mL bromophenol blue), boiled for 5min and analyzed by 10% SDS-PAGE gel for 2h at 100V. After that, gels were transferred to a nitrocellulose membrane for 1h at 100V in cold. Membrane were blocked with 5% nonfat milk and then incubated with the following antibodies: anti-IL-17RD, anti-NF-κB, anti-ERK and anti-p38 (Supplementary Table 4). Anti-GAPDH and anti-PARP-1 were used as loading control. The blots were revealed by incubation with the corresponding anti-rabbit, - mouse or goat fluor-coupled secondary antibody (LI-COR Biosciences) for 1h at RT. After washing, the membranes were scanned with the Odyssey infrared imaging system (LI-COR, Biosciences) at a wavelength of 700–800 nm. Densitometric analysis was performed using ImageJ software.

### Evaluation of mRNA expression levels

Samples were lysed with Tri-Reagent (Sigma) and total RNA was obtained following manufacturer’s instructions. RNA concentration and quality was determined by spectrophotometry according to absorbance at 260nm and 260/280 ratio. Complementary DNA (cDNA) was synthesized using M-MLV retrotranscriptase kit (M1701, Promega) and random primers (B070-40, Biodynamics). Then, real-time qPCR was performed using 50ng of cDNA per reaction using TaqMan Universal Master Mix II, no UNG (Applied Biosystems) and Taqman probes (Supplementary Table 5, Applied Biosystems) targeting specifics transcripts of interest. Amplification of 18S was used as endogenous control (Taqman 18S control Reagent, Applied Biosystems). Determinations were performed on pooled samples containing proportional mix of cDNA samples from 2-3 culture replicates in the case of cell lines or from tumors from 3-5 different mice. Reactions for both the transcripts of interest and 18S were carried out in triplicate. A StepOnePlus Real-Time PCR System thermal cycler (Applied Biosystems) was used according to the manufacturer’s instructions. The results were analyzed by calculation comparative CT (ΔΔCT) method using the StepOne software. Finally, the formula fold change = 2^-ΔΔCT^ was used to calculate the relative expression of RNA.

### RNA Sequencing

After medium removal, cells were collected and lysed with TCL buffer (Qiagen) with 1 % of β-mercaptoethanol and stored at −80°C until subsequent analysis. RNA was isolated using a Single Cell RNA purification kit (Norgen), including RNase-Free DNase Set (Qiagen) treatment. The RNA integrity number was evaluated with an Agilent RNA 6000 pico kit (all RIN>8). Retrotranscription was carried out with SMART-Seq v4 Ultra Low Input RNA Kit for Sequencing. Barcoded Illumina compatible libraries were generated from 5 to 10ng cDNA of each sample, using Nextera XTP reparation Kit. Libraries were sequenced on an Illumina Novaseq 6000 using 100bp paired-end mode, 20 million reads per sample.

### RNA-sequencing analysis

Quality profiles of sequencing files were evaluated using the FASTQC tool (“Babraham Bioinformatics - FastQC A Quality Control tool for High Throughput Sequence Data,” n.d.). Reads were filtered to remove adapters and low-quality bases using Trim_galore v0.6.5 (“Babraham Bioinformatics - Trim Galore!,” n.d.). The filtered sequences were mapped against the murine reference genome (ENSEMBL: mm10.GRCm38.97) using Subread v2.0.1 (36). Quantification of the mapped profiles was performed using the featureCounts v1.6.4 tool (37). Normalization of the quantified profiles and differential gene expression analysis was performed using the Bioconductor edgeR v3.28.0 package (38). Genes with a count greater than 10 in at least 70 % of the samples were kept (39). Then, samples were normalized using the TMM (for trimmed mean of M-values) method (40) to minimize differences in RNA composition. Differential gene expression was evaluated by fitting a negative binomial generalized linear model and differential expression was determined using the quasi-likelihood F-test (41). For further bioinformatics analysis genes were filtered by p value < 0.05 and log2fold change ≥ |1.5|. Heat maps were design using the excel spreadsheet and the clustvis tool (https://biit.cs.ut.ee/clustvis/), gene grouping was performed using the “Gene list analysis” Panther tool (http://pantherdb.org/). Volcano plot was performed in the R environment using the EnhancedVolcano package. Pathway analysis were performed using “Ingenuity Pathway Analysis” (QIAGEN) software. Data were stored in an NIH repository (https://www.ncbi.nlm.nih.gov/sra) under accession number PRJNA855854.

### Analysis of publicly available human tumor datasets

Data corresponding to the expression level of IL-17RA, IL-17RC and IL-17D transcripts in human tumor samples were obtained from cBioPortal for Cancer Genomics (https://www.cbioportal.org/). This online tool allows the visualization, analysis and download of tumor genomics datasets from multiple databases. For the present work, mRNA expression data of transcripts from 34 RNA-seq datasets of cancer studies belonging to the TCGA database were downloaded in November 2021. The data were processed in Phyton environment.

Overall survival analysis (defined as time to death), harmonized RNA-seq bulk data provided by TCGA, obtained from 472 melanoma samples (TCGA-SKCM) or 151 acute myeloid leukemia samples (TCGA-LAML), were used. The median gene expression level of IL-17RA, IL-17RC or IL-17RD were used as a cut-off point to segregate cancer patients into two groups: low and high expression. Then, Kaplan-Meier curves were generated for each tumor type stratified according to the expression level of IL-17RA, IL-17RC and IL-17RD in the aforementioned groups. The statistical significance of the overall survival curves was determined by a Log-rank test (R survival package).

### Statistical analysis

Statistical analysis was performed using GraphPad Prism 7.0 software. Several tests were performed including t-test, one-way ANOVA, two-way ANOVA or Kruskal-Wallis test as indicated in the legend of each figure. P values < 0.05 were considered significant.

## RESULTS

### Defects in the expression of IL-17RA or IL-17A and IL-17F in murine hosts lead to a disparate progression in different syngeneic tumor models

As a first step to understand the biological relevance of host IL-17 signaling in tumor progression, we aimed at identifying syngeneic tumor models that showed different outcomes of tumor progression in mice deficient for IL-17RA (IL-17RA KO mice) or IL-17A and IL-17F (IL-17A/F DKO mice) expression. To this end, we injected cell lines that give rise to different types of cancer including B16.SIY (melanoma), MC57.SIY (fibrosarcoma), EL4.SIY (lymphoma), and C1498.SIY (acute myeloid leukemia), and evaluated their growth in both deficient mice in comparison to wild type (WT) mice. We observed that the MC57.SIY fibrosarcoma, a tumor model that is completely rejected around day 12 post-injection (pi) in immunocompetent hosts, showed increased tumor volume in IL-17RA KO mice compared to WT controls at day 6 and 8 pi, though it is finally eliminated in both mouse strains (Figure 1A, left panel). Alike, B16.SIY melanoma growth was significantly increased in IL-17RA KO compared to WT mice at several day pi (Figure 1A, right panel). In contrast, the EL4.SIY lymphoma had a significant delayed progression in IL-17RA KO mice when compared to WT hosts (Figure 1B, left panel). Also, the tumors generated by the subcutaneous injection of C1498.SIY leukemia cells showed significant reduced volumes in mice lacking IL-17RA expression (Figure 1B, right panel). Similar outcomes in tumor progression were obtained using IL-17A/F DKO mice which, in comparison to WT mice, showed increased MC57.SIY and B16.SIY tumor volume on the peak of tumor growth or at the endpoint, respectively (Figure 1C); and reduced EL4.SIY and C1498.SIY tumor growth at endpoint (Figure 1D). These results illustrate that IL-17 signaling in the hosts has a significant impact on tumor growth, with contrasting (promotive or suppressive) effects depending on the tumor type, a notion that is in line with published evidence.

**Figure 1:**
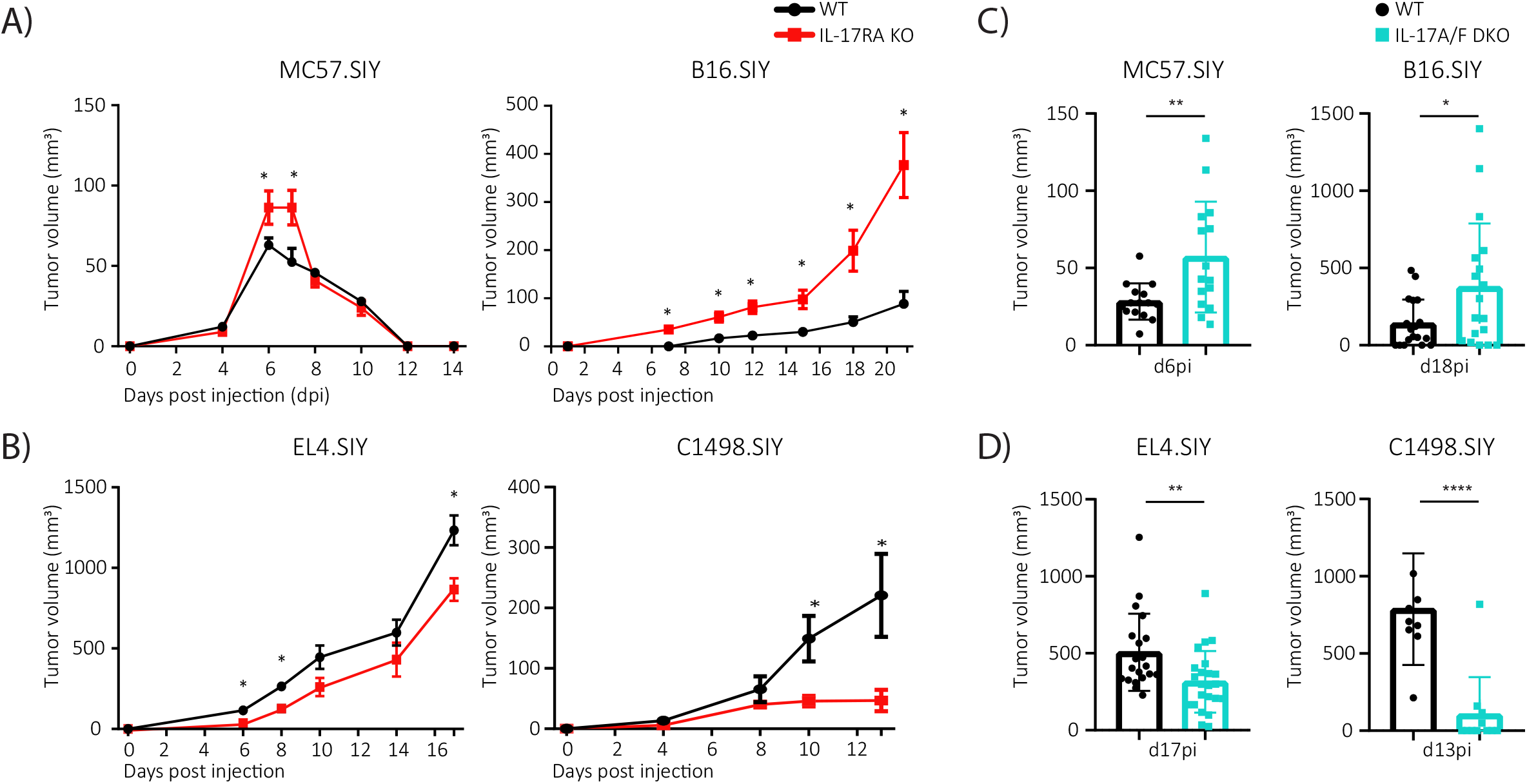
Deficiency in host IL-17 signaling promotes or suppresses tumor growth depending on tumor type. MC57.SIY, B16.SIY, EL4.SIY or C1498.SIY cells were injected subcutaneously into the right flank of WT, IL-17RA KO and IL-17A/F DKO C57BL/6 mice. **(A, B)** Curves of average tumor volume measured at different day post injection (dpi) of the indicated tumor cells in IL-17RA KO mice (red) and WT controls (black). **(C, D)** Tumor volumes at the peak of MC57.SIY tumor progression (d6pi) and at endpoint for B16.SIY, EL4.SIY and C1498.SIY (days 18, 17 and 13 pi, respectively) developed in IL-17 A/F DKO (blue) and WT mice (black). In **A-D** results are shown as mean ± SD. N=10-20 from 2 or 3 independent experiments. In **A** and **B** P-values were calculated by multiple t-test. In **C** and **D** P-values were calculated by unpaired t-test. * p < 0.05; ** p < 0.01; *** p < 0.001; **** p < 0.0001.

### Lack of IL-17RA or IL-17A and IL-17F expression in host cells diversely impacts the expression of inflammatory and angiogenic mediators in different tumors

Within the tumor microenvironment, IL-17 signaling may impact on different cell types modulating the expression of proteins able to regulate inflammation and angiogenesis, two tightly linked events critical for tumor progression. Thus, we aimed to evaluate the expression of different proteins involved in these processes as an initial step to elucidate the causes of distinctive tumor growth in conditions of defective IL-17 signaling in murine hosts, selecting B16.SIY melanoma and EL4.SIY lymphoma as models of contrasting progression outcomes. Using a proteome profiler antibody array, we interrogated the expression of 53 proteins and found that 27 were expressed in lysates from both tumor types developed in WT, IL-17RA KO and IL-17A/F DKO mice (Figure 2A and Supplementary Figure 1A). Additionally, 11 to 14 of the proteins detected were significantly over or under-expressed in tumors from IL-17RA KO and/or DKO mice, in comparison to the corresponding tumor from WT counterparts (Supplementary Figure 1B and C). Notably, the impact of a deficient host IL-17 signaling in the expression of inflammatory and angiogenic proteins were essentially different in B16.SIY and EL4.SIY tumors (Figure 2B). In fact, very few changes in protein expression such as the downregulation of IGFBP-3, MMP-3, MMP-9 and Serpin E1 and the upregulation of IGFBP-2, were shared between B16.SIY and EL4.SIY tumors from IL-17RA KO and IL-17A/F DKO, respectively. To identify particular proteins linking IL-17 signaling and tumor progression in our models, we focused on mediators that were up- or down-regulated in at least a two-fold magnitude (log2(FC)| ≥ 1) in tumors from IL-17RAKO or IL-17A/F DKO mice, compared to tumors from WT controls (gray areas in Figure 2C). We found that, IGFBP-2, a protein that promotes several key oncogenic processes (42), exhibited the highest expression in B16.SIY tumors from IL17RAKO mice which, in turn, presented increased growth when compared to tumors from WT controls. Differently, the reduced size of EL4.SIY tumors in IL-17RA KO mice correlated with a marked down-regulation of PTX3, a tumor-derived protein that promotes migration, invasion, and/or epithelial to mesenchymal transition (43, 44). In regard to IL-17A/F DKO hosts, increased growth of B16.SIY tumors were associated with a notable downregulation of MMP-3 and MMP-9 expression which are a direct target of IL-17 (45). In contrast, the reduced progression of EL4.SIY tumors occurred concomitantly with a robust up-regulation of Serpin F1, a protein with anti-angiogenic, anti-tumorigenic and anti-metastatic properties (46). Overall, this exploratory evaluation fosters the idea that the tumor type may be a critical determinant of the responsiveness to IL-17 by the cells of the tumor microenvironment.

**Figure 2.**
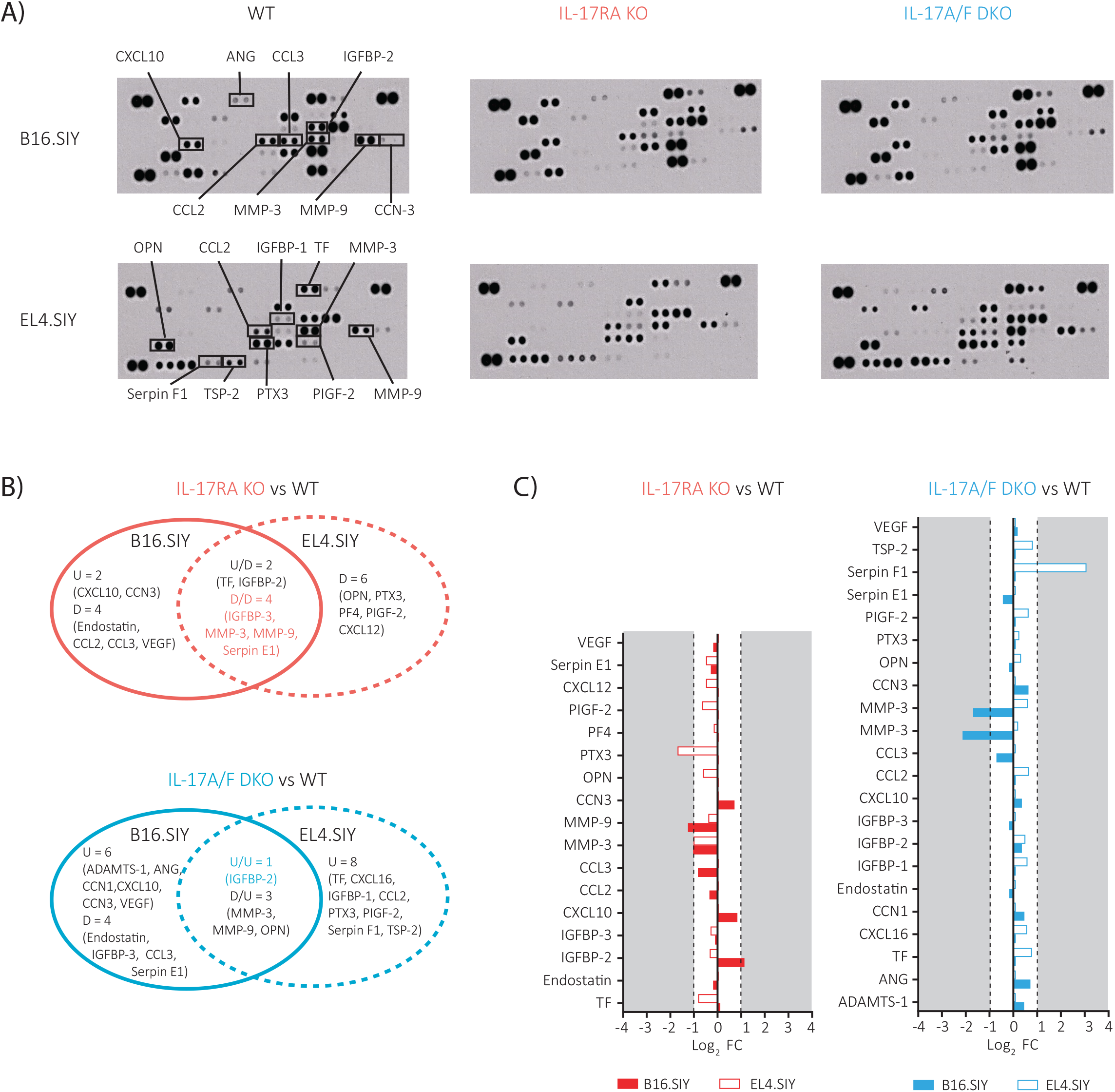
Defective host IL-17 signaling differently impacts the expression of tumor inflammatory and angiogenic proteins depending on tumor type. **(A)** Proteome Profiler Mouse Angiogenesis Array Kit (ARY015, R&D) was used to assess the relative levels of inflammation- and angiogenesis-related proteins in the B16.SIY and EL4.SIY tumors lysates collected at 17-18 dpi from WT, IL-17RA KO and IL-17A/F DKO mice. Selected proteins of interest are annotated. **B-C)** Differentially expressed proteins of tumors developed in IL-17RA KO (red) or IL-17A/F DKO mice compared to WT mice (black). **(B)** Venn diagrams show proteins that are differentially expressed in B16.SIY versus EL4.SIY tumor lysates developed in IL-17RA KO (red) compared to WT mice (black) (upper diagram) and IL-17A/F DKO mice (blue) compared to WT mice (black) (bottom diagram). The direction of change is indicated (U=upregulation and D=downregulation). Proteins with changes in similar direction between B16.SIY and EL4.SIY tumor lysates are shown in color. **(C)** Bar graphs the magnitude and direction of changes of the between B16.SIY (filled bars) and EL4.SIY (empty bars) tumor lysates. Protein levels are expressed as log2 fold change (FC), and gray areas indicate |log2 FC| values > 1.

### Defective host IL-17/IL-17RA pathway compromises antitumor CD8+ T cell immunity irrespective of tumor type and progression

Considering that the immune component may play a pro- and anti-tumor role and that IL-17 may modulate its function, we performed an evaluation of the tumor immune infiltrate in B16.SIY and EL4.SIY models. We initially compared immune infiltrates in tumors developed in WT hosts and determined that the melanoma model contained significantly higher frequency of CD45+ immune cells (Figure 3A). To define the distribution of the main innate and adaptive immune cell subsets present in the tumor infiltrate, we performed multiparametric flow cytometry with a 16-marker panel. Unsupervised analysis using the dimension reduction algorithm Uniform Manifold Approximation and Projection (UMAP) followed by FlowSOM clustering algorithm led to the identification of ten clusters (P0-P9) (Figure 3B) which were annotated according to the relative intensity of expression of different population markers (Supplementary Figure 2A). B16.SIY and EL4.SIY tumors showed differences in the frequency of certain subpopulations, in particular, neutrophil-like cells (P8), NK cells (P7) and CD4+ T cells (P3). These differences were confirmed by statistical analysis that showed that B16.SIY tumors contained significantly increased infiltration of a population with neutrophil markers (P8) and reduced presence of NK cells (P7) and CD4+ T cells (P3) in comparison to EL4.SIY tumors (Figure 3C).

**Figure 3:**
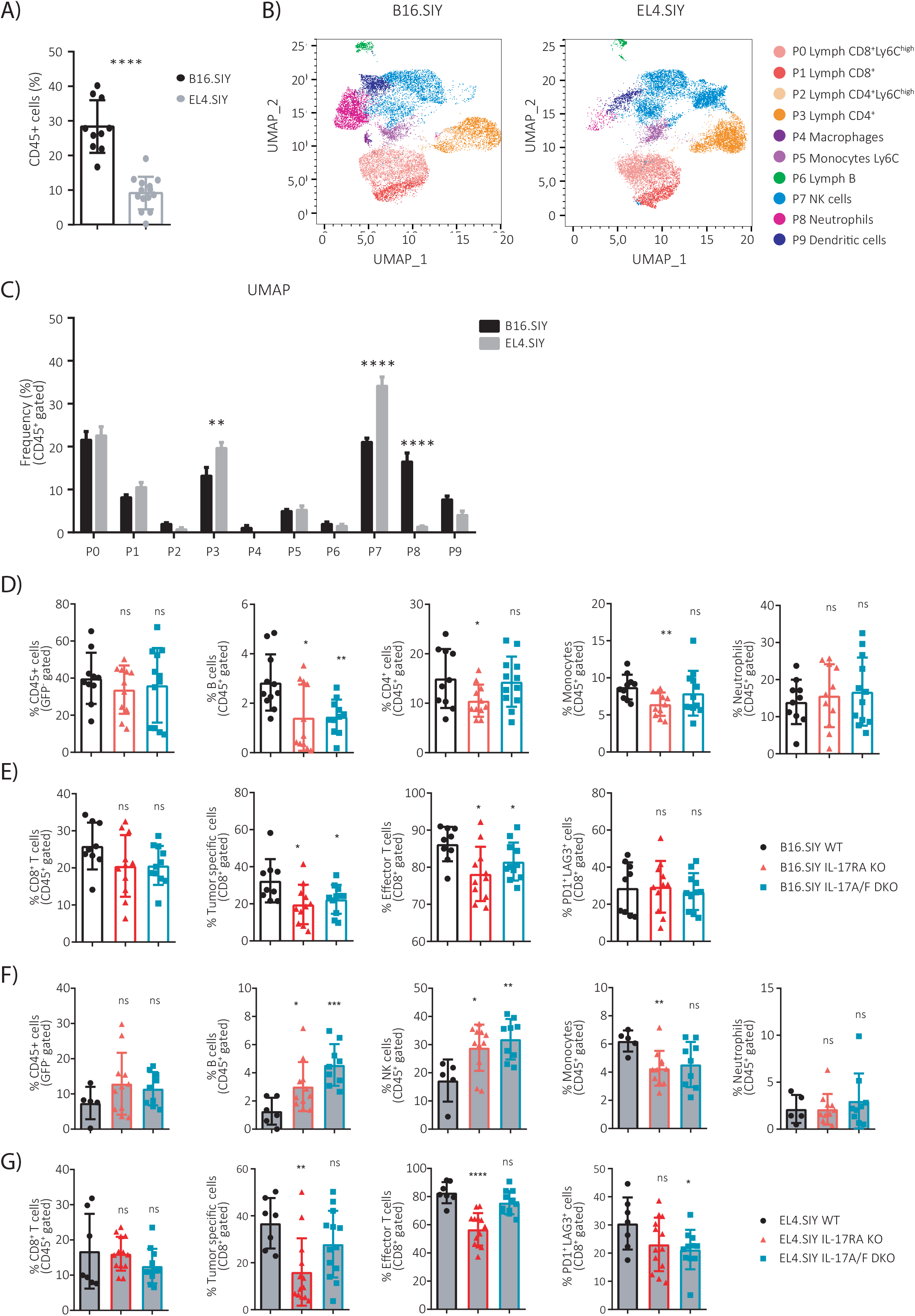
Deficiencies in host IL-17A/F:RA pathway compromise tumor-specific CD8+ T cell immunity in B16.SIY and EL4.SIY bearing mice and affect other tumor-infiltrating subpopulations in a dissimilar manner depending on tumor type. **(A-C)** Percentage of CD45+ cells within total live cells in B16.SIY and EL4.SIY tumors from WT mice collected at d18pi and d17pi, respectively. **(B)** Plots show the ten FlowSOM-identified clusters (P0-P9) from the CD45+ population overlaid on the UMAP projection in the B16.SIY and EL4.SIY tumors evaluated in A. **(C)** Comparison of the frequencies of each cluster defined in B in B16.SIY (black) and EL4.SIY (gray) tumors evaluated in A. **(D-G)** Frequency of the indicated immune subpopulations in B16.SIY (**D-E**, empty bars) and (**F-G**, filled bars) EL4 tumors collected at d18pi and d17pi, respectively from WT (black), IL-17RA KO (red) and IL-17A/F DKO (blue) mice. P-values were obtained by two-tailed Student’s t-test. * p < 0.05; ** p < 0.01; *** p < 0.001; **** p < 0.0001.

Afterwards, we focused on understanding how lack of IL-17RA or IL-17A and IL-17F expression in the host impacted on the immune cell infiltrate composition in each tumor type. To this end, we evaluated the main immune subsets present in B16.SIY and EL4.SIY tumors from IL-17RAKO and IL17A/F DKO mice in comparison to those from WT mice using the same panel as mentioned before. Considering that IL-17 signaling has been associated with more robust (47) as well as exhausted (48) CD8+ T cell responses, we also compared the specificity and the activation and exhaustion phenotype of these cells in each strain/tumor combination. The lack of IL-17RA or IL-17A/F expression in murine host did not affect the extent of infiltration in B16.SIY tumors as illustrated by the conserved frequency of CD45+ cells in comparison to B16.SIY tumor from WT mice (Figure 3D, left panel). However, the distribution of major immune cell subsets showed significant changes that included the reduction of infiltrating B cells in B16.SIY tumors from IL-17RA KO and IL-17A/F DKO mice as well as decreases in CD4+ T cell and monocyte frequencies in tumors from IL-17RA KO mice (Figure 3D). No changes were observed in the frequency of neutrophils (Figure 3D). Additionally, mice with deficient IL-17 signaling exhibited a conserved infiltration of total CD8+ T cells in B16.SIY tumors (Figure 3E, left panels) but reduced frequencies of tumor-specific (defined according to their staining with a SIY-dextramer) and effector CD44^high^CD62L^low^ CD8+ T cells (Figure 3E, middle panels). Frequency of tumor exhausted-like (PD1+Lag3+) CD8+ T cells were comparable in the three experimental groups (Figure 3E, right panels). As observed in B16.SIY tumors, host deficiencies in IL-17RA and IL-17A/F expression did not affect the magnitude of the CD45+ cell infiltration in EL4.SIY tumors (Figure 3F, left panel). However, EL4.SIY tumor infiltrates from IL-17RA KO and DKO mice contained higher frequencies of B cells and NK cells in comparison to WT counterparts (Figure 3F, middle panels). Monocytes were also reduced in EL4.SIY tumors from IL-17RA KO mice while neutrophils showed similar frequencies in tumor from the three mouse strains (Figure 3F, right panels). Total CD8+ T cells were found at similar frequencies in EL4.SIY tumors from WT, IL-17RA KO and IL-17A/F DKO (Figure 3G, left panel). Nevertheless, EL4.SIY tumors in IL-17RA KO mice presented reduced infiltration of tumor specific and effector CD8+ T cells (Figure 3G, middle panels) and conserved infiltration of exhausted-like CD8+ T cells (Figure 3G, right panels). In contrast, IL-17A/F DKO mice developed EL4.SIY tumors that exhibited conserved percentages of tumor specific and effector CD8+ T cells (Figure 3G, middle panels) and reduced frequencies of exhausted CD8+ T cells (Figure 3G, right panels). The contrasting findings in the tumor-specific CD8+ T cell responses to EL4.SIY in IL-17RA KO and IL-17A/F DKO mice may result from the fact that host IL-17 signaling is completely impaired by lack of IL-17RA expression in IL-17RA KO mice but not in IL-17A/F DKO mice where lack of endogenous IL-17 could be, at least partially, compensated by IL-17 production by the EL4.SIY tumor as we will show in the following section.

Altogether, our results about how host IL-17 signaling modulate immune cell infiltrates in B16.SIY and EL4.SIY tumors reinforced the notion of a tumor-type dependency in these biological effects. Our data also underlined that, when host IL-17 signaling is impaired, the tumor-specific CD8+ T cell immunity is reduced, but this reduction does not necessarily correlate with enhanced tumor progression.

### Tumor cell lines exhibit distinct expression patterns of IL-17 cytokines and receptors that are associated with particular IL-17 gene signatures

Given that tumor cells are the main cell population within the tumor, whether they differentially express cytokines and receptors of the IL-17 family may determine the overall effect of IL-17 in tumor progression. To test this idea, we evaluated the expression of cytokines and receptor subunits belonging to the IL-17A/IL-17RA pathways in the tumor cell lines used in Figure 1. By real-time PCR, we observed that *Il17a* transcripts were only detected in EL4.SIY and C1498.SIY cells, while *Il17f* mRNA was present in all four cell lines evaluated (Figure 4A). Regarding the expression of the IL-17R subunits, we determined that *Il17ra* and *Il17rd* transcripts were detected in all the cell lines while presence of *Il17rc* mRNA was limited to MC57.SIY and B16.SIY cells (Figure 4B). To further validate these results at protein level, we evaluated the concentration of IL-17A and IL-17F in supernatants obtained after culturing the different cell lines. As depicted in Figure 4C, IL-17 production was limited to EL4.SIY and C1498.SIY cells in concordance with the presence of transcripts encoding these cytokines in both cell lines. Remarkably, IL-17F was detected only in the supernatants of EL4.SIY and C1498.SIY despite the fact that *Il17f* mRNA was present in all the cell lines evaluated. In agreement with transcriptional data, expressions of IL-17RA and IL-17RD proteins were detected in the four cell lines, while IL-17RC was limited to MC57.SIY and B16.SIY cells, as determined by flow cytometry or western blot (Figure 4D and E). Altogether, these results highlight particular expression profiles of IL-17 cytokines and receptors in cell lines of different tissue origins. In sum, IL-17RA and IL-17RD were ubiquitously expressed whereas IL17RC expression was limited to tumor cell lines of epithelial/mesenchymal origin. On the other hand, IL-17A and IL-17F secretion was limited to hematopoietic cell lines.

**Figure 4:**
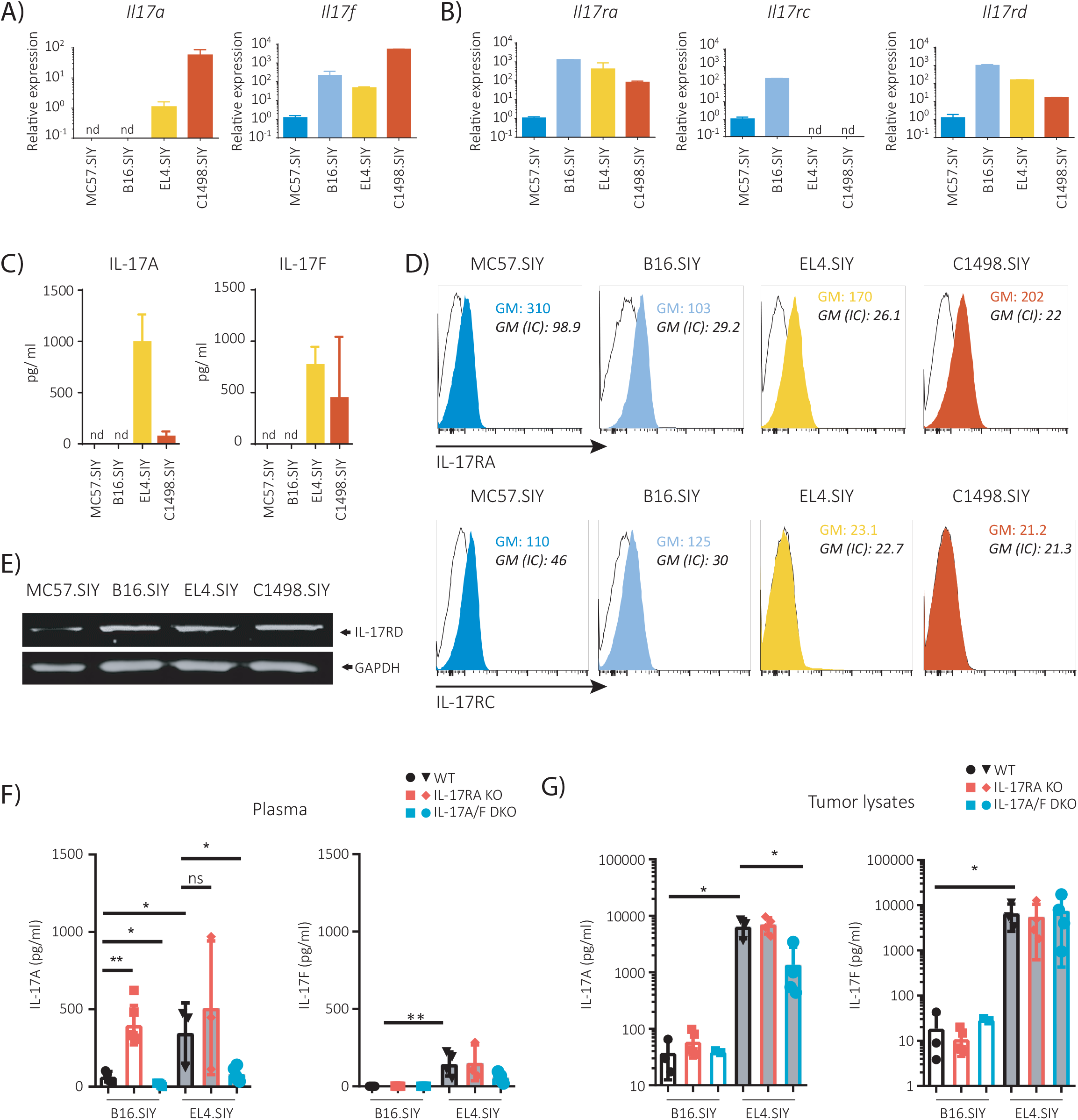
Cell lines and tumors of different tissue origins show particular expression profiles of receptors and cytokines of the IL-17A/F:RA pathway. **(A)** Relative amounts of the transcripts encoding IL-17A and IL-17F cytokines in B16.SIY (melanoma), MC57.SIY (fibrosarcoma), EL4.SIY (lymphoma) and C1498.SIY (acute myeloid leukemia) cells. **(B)** Relative amounts of *Il17ra, Il17rc* and *Il17rd* transcripts in the indicated tumor cell lines. **(C)** Concentration of IL-17A and IL-17F determined by ELISA in culture supernatants of the indicated tumor cell lines. **(D)** Representative histograms depicting IL-17RA and IL-17RC expression determined by flow cytometry in the indicated tumor cell line. Isotype control (IC) staining is shown in black histogram. **(E)** Expression of IL-17RD evaluated by western blot in lysates from the indicated tumor cell lines. **(F-G)** IL-17A and IL-17F concentration determined by ELISA in plasma **(F)** and tumor lysates **(G)** collected from WT (black), IL-17RA KO (red) or IL-17A/F DKO B16.SIY or EL4.SIY tumor-bearing mice at endpoint (d18pi and d17pi respectively). Transcript amounts in RT-qPCR were normalized to 18S control gene transcript level and then relativized to MC57 transcript levels, n=3 replicates per group. nd: non detectable. In **F** and **G**, N=3 mice/group, one-way ANOVA was used for statistical analysis: * p <0.05 **; p <0.01; *** p <0.001; ****p < 0.0001.

Next, we evaluated the systemic and local production of IL-17A and IL-17F in WT and IL-17 signaling deficient mice bearing B16.SIY and EL4.SIY tumors. Systemic levels of IL-17A were significantly higher in B16.SIY-bearing, but not in EL4.SIY-injected, IL-17RA KO mice compared to WT counterparts (Figure 4F, left panel). In addition, IL-17A concentrations in plasma of IL-17A/F DKO mice injected with B16.SIY or EL4.SIY cells were reduced compared to WT mice (Figure 4F, left panel). Moreover, plasma IL-17F concentrations were practically undetectable in all groups of mice injected with B16.SIY cells and were low but comparable between all the strains bearing EL4.SIY tumors (Figure 4F, right panel). Local levels of IL-17A and IL-17F in tumor lysates were similar in all the B16.SIY tumor-bearing mice (Figure 4G). In coherence with our previous *in vitro* studies, IL-17A and IL-17F were markedly more abundant in EL4.SIY tumors relative to B16.SIY tumors, and showed equivalent concentrations in tumors from all mouse strains. Of note, the only exception was IL-17A/F DKO mice, whose EL4.SIY tumor lysates showed significantly reduced amounts of IL-17A compared to WT (Figure 4G, left panel).

Given the results described above, we decided to follow up with a deep and unsupervised evaluation of the gene signature associated with the IL-17 pathway in selected cell lines by RNA sequencing. As expected, due to their different tissue origin, EL4.SIY and B16.SIY cells showed substantial differences in gene expression as illustrated by the 11694 differentially expressed genes (DEGs) (cut off: ļlog2(FC)≥ 1.5, p <0.05), of which 5774 transcripts were highly expressed on EL4.SIY cells and 5920 transcripts were more abundant in B16.SIY cells (Supplementary figure 3A). Analysis of the DEGs obtained by RNA-seq associated with the IL-17 family confirmed our data using RT-PCR regarding particular expression of molecules from the IL-17 pathway on tumor cells. In fact, *Il17rc* gene expression was higher in B16.SIY than in EL4.SIY cells, while *Il17a* and *Il17f* transcripts were more abundant in EL4.SIY cells. Besides, *Il17ra* and *Il17rd* genes were similarly present in both cell lines. Interestingly, examination of the 49 genes of the “IL-17 pathway” retrieved from the KEGG database revealed that 28 genes (57%) showed contrasting expression between B16.SIY and EL4.SIY cells. From these 28 DEGs, selected genes are highlighted in red in the volcano plot from Supplementary figure 3A, and the complete list is depicted in a clustered heatmap of Supplementary figure 3B. Altogether, these results underscored that cells from different tissue origin show different patterns of expression of cytokines and receptors of the IL-17A/IL-17RA pathway, paralleled with particular activation of the IL-17 related gene signature.

### *In vitro* stimulation with IL-17 induces distinct responses in different tumor cell lines

We next evaluated how *in vitro* stimulation with IL-17 affected tumor cell lines biology. We cultured the four different cell lines with IL-17A and IL-17F, and evaluated changes in proliferation, cell migration and secretion of inflammatory cytokines and chemokines. Using an MTT assay, we first established that stimulation with IL-17A and IL-17F during 24 h did not change the cell counts of the different tumor cell lines with the only exception of MC57.SIY cells, which showed reduced cell numbers upon culture in the presence of IL-17A (Supplementary Figure 4A). We also evaluated if IL-17A and IL-17F affected cell migration capacity of adherent cell lines (B16.SIY and MC57.SIY) through a wound healing assay, finding that IL-17A and IL-17 had no effect on cell migration up to 12 h of culture (Supplementary Figure 4B). Finally, we observed that addition of IL-17A and IL-17F to the tumor cell cultures induced the production of CXCL1 by the B16.SIY and MC57.SIY cells, but not by EL4.SIY and C1498.SIY (Figure 5A). This inducible effect was not observed for the secretion of IL-1β, IL-6, TNF or CXCL2 by any of the tumor cell lines (Supplementary Figure 4C).

**Figure 5.**
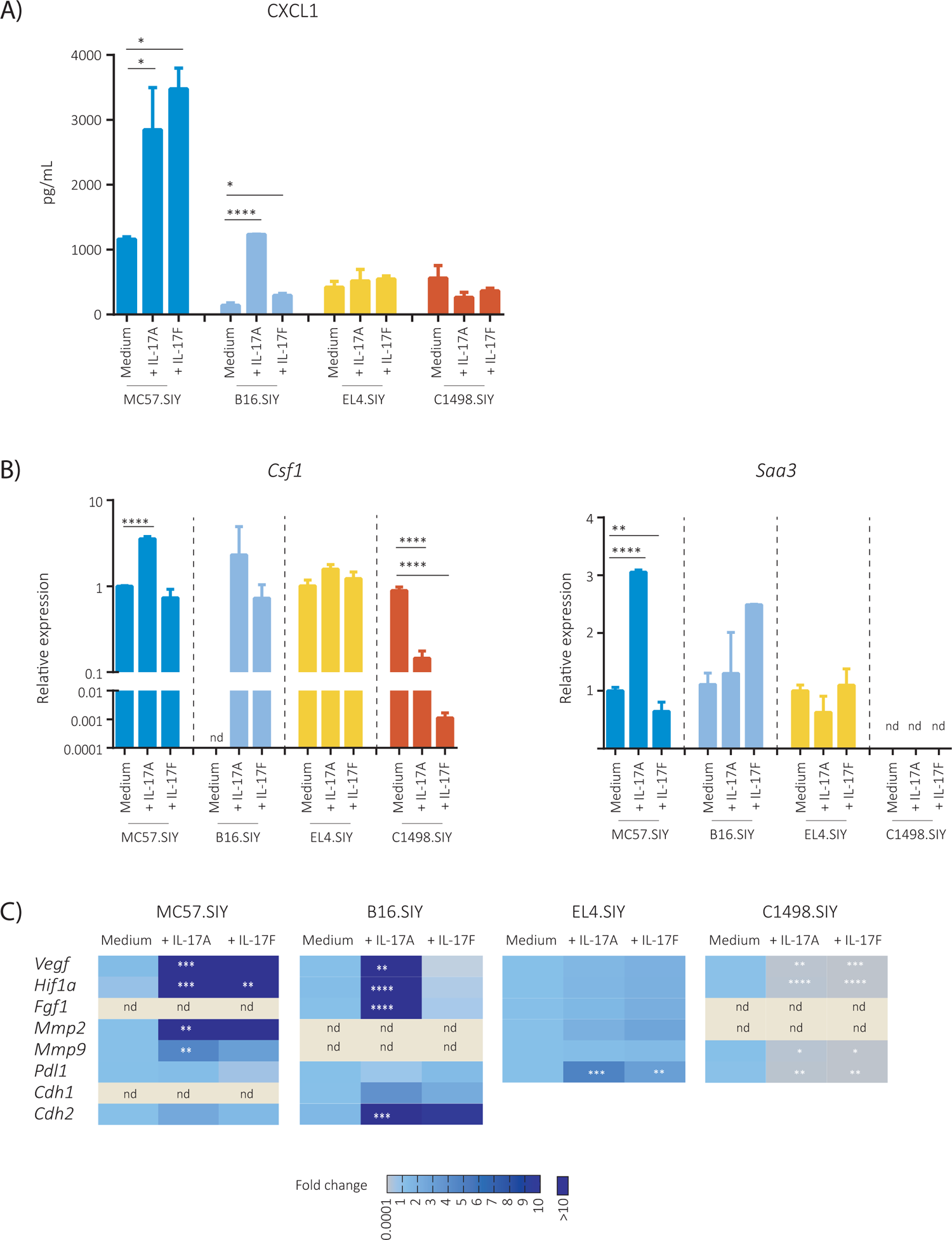
In vitro stimulation with IL-17A and IL-17F distinctly modulates the transcript level of target genes and the expression of mediators associated with tumor progression in different tumor types. B16.SIY, MC57.SIY, EL4.SIY and C1498.SIY cells were cultured in the presence of IL-17A or IL-17F (200ng/mL) in serum reduced conditions. **(A)** Concentration of CXCL1 was determined by ELISA in supernatants after 48h of stimulation. **(B)** Relative amounts of Csf1 and Saa3 mRNA determined by RT-qPCR after 24h of stimulation. **(C)** Heat-map summarizing relative levels of *Hif1a, Vegf, Mmp2, Mmp9, Fgf, Pdl1, E-cadherin* and *N-cadherin* transcripts determined by RT-qPCR after 24 h of stimulation. Transcript amounts were normalized to 18S. nd: non detectable. One-way ANOVA was used for statistical analysis: *p value <0.05; **p value<0.01; ***p value<0.001; ****p value< 0.0001.

Subsequently, we focused on the transcriptional evaluation of genes conventionally associated to an IL-17 gene signature such as those encoding GM-CSF and Serum Amyloid 3 proteins (2). We showed that the stimulation of B16.SIY and MC57.SIY cell lines with IL-17A increased the amounts of *Csf1* and/or *Saa3* mRNA while stimulation of B16 with IL-17F augmented the levels of *Csf1*. In contrast, stimulation of EL4.SIY and C1498.SIY with IL-17A or IL-17F had no effect or reduced the amounts of these gene transcripts (Figure 5B). We also quantified other transcripts that encoded different proteins reported to be modulated by IL-17 and considered relevant for cancer biology or antitumor immunity (49–51). As depicted in the heat map from Figure 5C, stimulation with IL-17A and/or IL-17F significantly induced the transcription of *Vegf, Hif1a, Mmp2, Mmp9, Cdh1* and/or *Chd2* in B16.SIY and MC57.SIY but showed no effect, or even inhibited, the transcription of these genes in EL4.SIY and C1498.SIY cell lines. Notably, a different pattern of response was observed for the *Pdl1* gene, that remained unchanged in B16.SIY and MC57.SIY cell lines upon stimulation with IL-17A and IL-17F but was significantly induced in EL4.SIY cells and inhibited in C1498.SIY cells. Overall, our data highlight that IL17A and IL-17F stimulation triggers particular responses in cell lines with specific IL-17R subunit expression patterns.

### Tumor cell lines with different expression pattern of IL-17R subunits show particular cell signaling and gene signatures downstream IL-17 stimulation

Canonical signaling downstream IL-17RA drives the activation of MAPK and the classical NF-κB pathway that results in a gene signature associated with inflammation. However, many studies have highlighted celltype specific responses to IL-17A that may be consequence of the activation of non-canonical signaling and a diverse gene expression program (2). Considering these antecedents, we selected B16.SIY and EL4.SIY cell lines and evaluated the intracellular signaling and gene transcriptional program triggered after IL-17A stimulation. We first focused on the activation of the MAPKs and determined that B16.SIY and EL4.SIY showed a basal phosphorylation of ERK and p38 that was not further increased by IL-17A at any time point evaluated (Supplementary Figure 5). Differently, we observed that phosphorylation of the p65 NF-κB subunit was almost undetectable in B16.SIY cells at basal level, but it was induced after 15-30 min of IL-17A stimulation in a weak but prolonged fashion (Figure 6A, left upper panel). Accordingly, IL-17A stimulation of B16.SIY cells during 30 min resulted in increased translocation of p65 to the nucleus (Figure 6A, right upper panel). Otherwise, EL4.SIY cells exhibited a basal phosphorylation and nuclear translocation of p65 that were not further increased by IL-17A stimulation (Figure 6A, left and right lower panels). Analyzed together, these data are consistent with the notion that IL-17A is only a modest activator of NF-κB (1), and underline that activation of the IL-17A mediated canonical signaling pathway may be cell-type specific and likely dictated by the expression pattern of IL-17R subunits.

**Figure 6.**
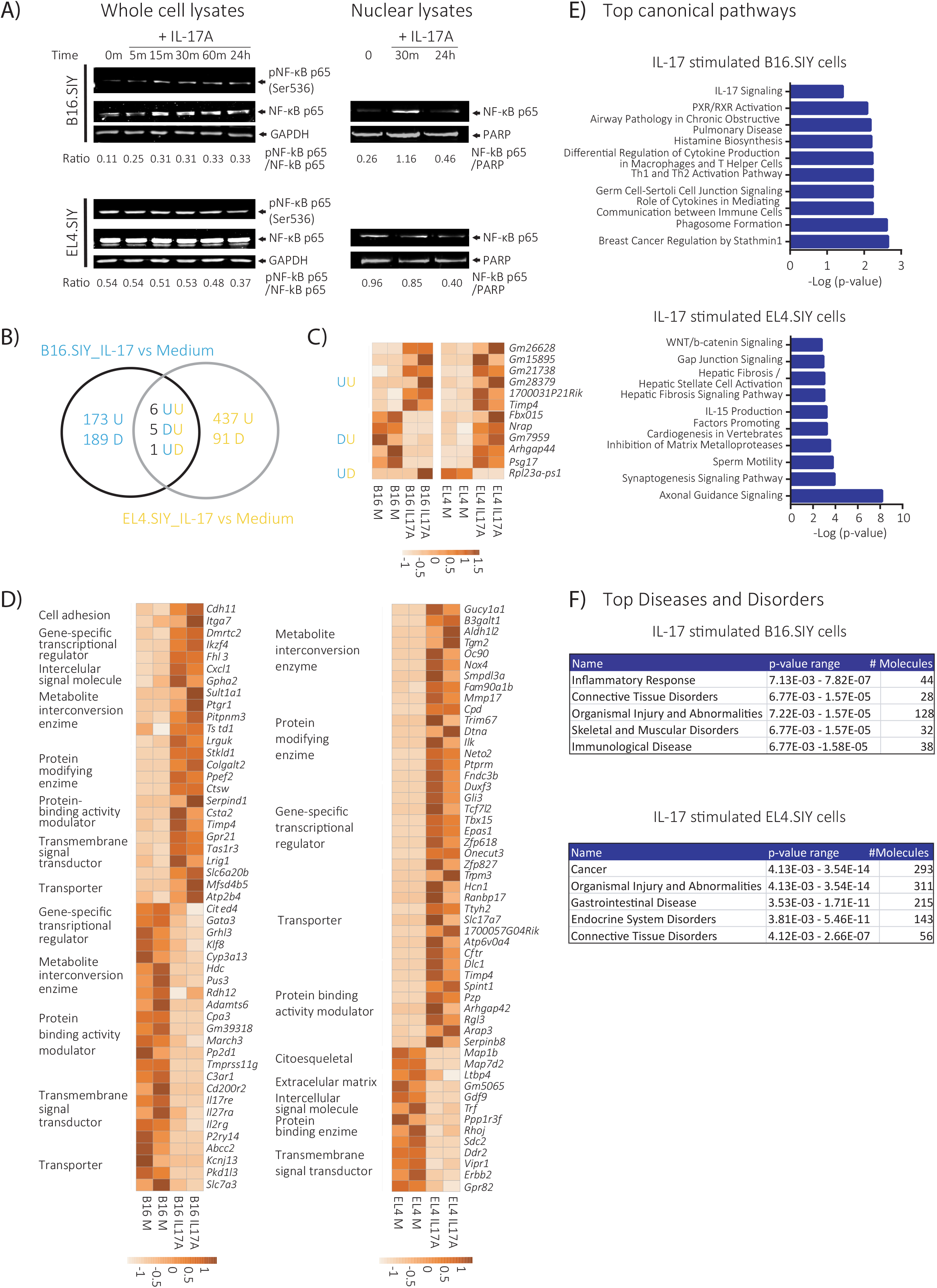
IL-17A activates particular signaling pathways and transcriptional signatures in B16.SIY versus EL4.SIY tumor cells. **(A)** Expression of phosphorylated (pNF-κB p65) and total NF-κB p65 subunit evaluated by western blot in whole cell lysates and nuclear lysates of B16.SIY and EL4.SIY after 0, 5, 15, 30, 60 minutes (m) and/or 24 h of stimulation with IL-17A (200 ng/mL) as indicated. GADPH and PARP expression were used as loading control. The indicated ratio values between the relative expression of the indicated proteins were determined using the FIJI software. **(B-F)** Analysis of RNA-seq performed in B16.SIY and EL4.SIY cells stimulated or not with IL-17A (100 ng/mL) for 24 h. **(B)** Venn diagram shows the comparison between the genes differentially modulated by IL-17 in B16.SIY (blue) and EL4.SIY (yellow) using a cut-off (|Log2FC|≥ 1.5, p-value < 0.05). The direction of the change is indicated as U: upregulation and D: downregulation. **(C)** Heat map for the expression of the 12 transcripts differentially modulated by IL-17A in both B16 and EL4 cells. **(D)** Heat maps of the top 50 genes modulated by IL-17 exclusively in B16.SIY or in EL4.SIY, grouped into families according to their biological function. **(E)** “Main canonical pathways” enriched in B16.SIY and EL4.SIY cells after IL-17A stimulation as identified by IPA. **(F)** “Major diseases and disorders” associated with IL-17A stimulation in B16.SIY and EL4.SIY cells as identified by IPA.

To gain insights into the global gene expression program triggered by IL-17A in B16.SIY versus EL4.SIY cells, we next evaluated their transcriptional profiles by RNA sequencing. We determined that 24 h stimulation with IL-17A resulted in 374 DEGs in B16.SIY cells and 540 DEGs in EL4.SIY cells compared to non-stimulated cells. (Figure 6B). Within these DEGs, we identified only 12 genes that were commonly modulated in both cell lines, 6 of which were upregulated in both cell lines, while the rest showed opposite responses. The other DEGs induced upon IL-17A stimulation involved 173 genes upregulated and 189 genes downregulated exclusively in B16.SIY cells, and 437 genes upregulated and 91 genes downregulated exclusively in EL4.SIY cells. Among the common 12 DEGs, there were several gene products involved in cellular organization and localization (Figure 6C). Differently, the genes that were significantly modulated by IL-17A exclusively in each cell line belonged to different gene categories (Figure 6D). Ingenuity Pathway Analysis (IPA) revealed that, within the Top Canonical Pathways enriched in IL-17A-stimulated B16.SIY cells, there were several pathways associated with cytokine-mediated responses such as “Role of Cytokines in Mediating Communication between Immune Cells” and “Th1 and Th2 Activation Pathway”, with the pathway “IL-17 signaling” also significantly enriched (Figure 6E, upper panel). In contrast, within the Top Canonical Pathways enriched in IL-17A-stimulated EL4.SIY cells there were several associated to signaling during cellular processes linked to cell growth and cell-to-cell interactions like “Axonal Guidance Signaling” and “Synaptogenesis Signaling Pathway” (Figure 6E, lower panel). Additionally, “Inflammatory Response” was the top identified “Diseases and Biofunctions” in IL-17A-stimulated B16.SIY cells, whereas “Cancer” appeared as the topmost one enriched in 17A-stimulated EL4.SIY cells (Figure 6F). Collectively, these data indicate that IL-17A induces significant transcriptional changes in both tumor cell lines but its effects are markedly divergent. Thus, B16.SIY cells that express IL-17RD as well as IL-17RA and IL-17RC which form the canonical IL-17R show a response to IL-17A that includes the activation of inflammatory pathways while EL4.SIY cells that express only IL-17RA and IL-17RD exhibit responses related to the induction of processes related to cellular biology.

### Blockade of IL-17RC changes B16.SIY cells response profile to IL-17 stimulation

The results described above support the notion that expression of IL-17RC may be a key determinant in the responsiveness of tumor cells to IL-17. To address this, we repeated the IL-17A stimulation experiments in B16.SIY cells adding a blocking IL-17RC antibody. We observed that IL-17RC blockade completely prevented the upregulation of the classical IL-17 target gene *Cfs1* in IL-17A-stimulated B16.SIY cells (Figure 7A), as well as, reduced the secretion of CXCL1 (Figure 7B). In contrast, blockade of IL-17RC resulted in a marked increase of *Pdl1* transcript levels (Figure 7C). The integrated analysis of the behavior of the three IL-17A targets shows that, in conditions of IL-17RC blockade, B16.SIY cells response to IL-17A resembles that of EL4.SIY cells. These findings indicate that IL-17RC signaling is strictly required for the induction of classical IL-17A target genes and proteins in B16.SIY cells. Interestingly, blockade of IL-17RC did not abrogate responsiveness to IL-17A but rather deviated the response to the upregulation of a different set of genes likely as a consequence of signaling through IL-17RA and IL-17RD.

**Figure 7.**
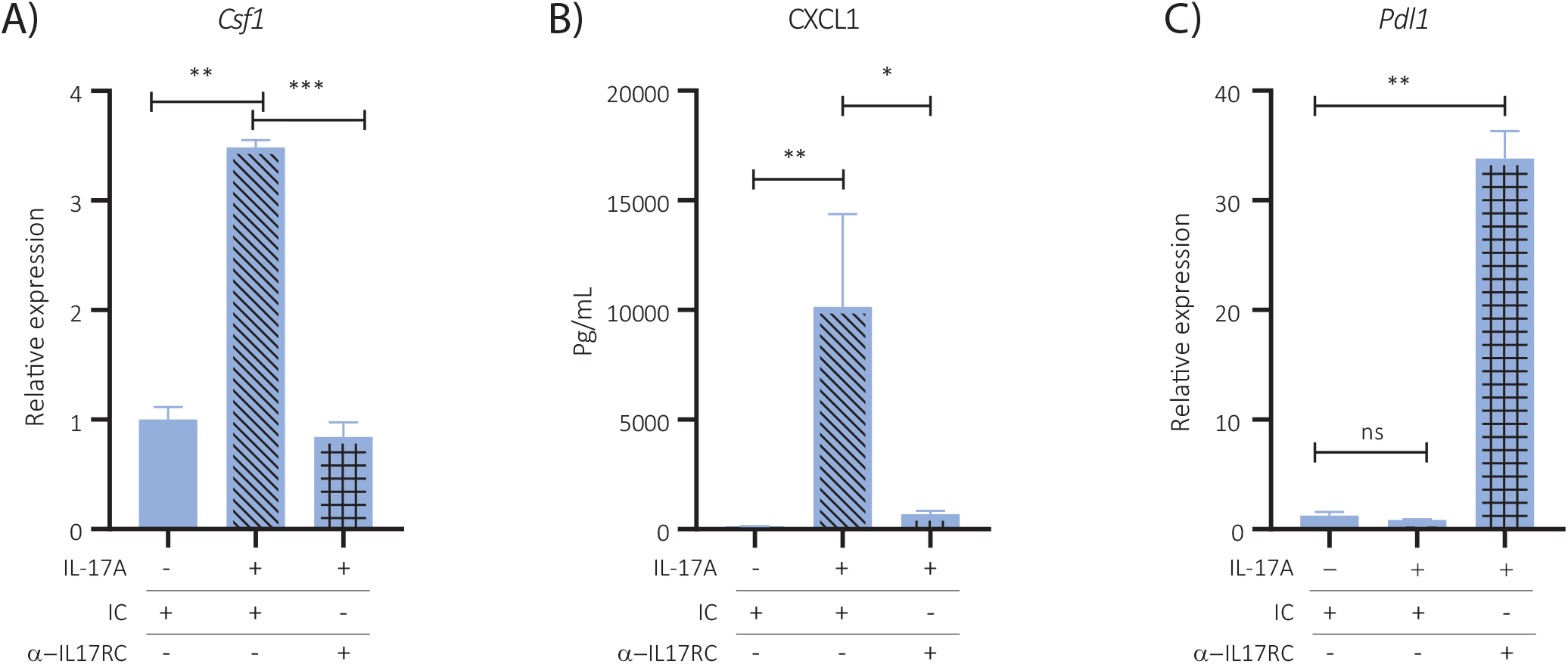
IL-17RC blockade modifies the response profile to IL-17A stimulation in B16.SIY cells. B16.SIY cells were stimulated during 24h under serum-reduced conditions with or without IL-17A (200ng) in the presence of an IL-17RC blocking antibody (5ug/mL) or an isotype control (IC). **(A)** Relative amounts of *Csf1* mRNA determined by RT-qPCR. **(B)** Concentration of CXCL1 determined by ELISA. **(C)** Relative amounts of *Pdl1* transcripts determined by RT-qPCR. Levels of *Csf1* and *Pdl1* transcripts were relativized to the levels ofIC-treated samples. Results are shown as the mean ± SD of 3 replicates for each condition. P-values were calculated by Two-tailed unpaired t-test. Ns: non-significant.

### Human cancers show different expression profiles of IL-17R subunits that correlate with different clinical outcomes

To investigate the clinical relevance of our findings, we examined the expression of IL-17R subunits in human tumors by analyzing data from The Cancer Genome Atlas (TCGA). Significant heterogeneity in *il17ra, il17rc* and *il17rd* transcript expression was observed across different cancer types (Supplementary Figure 6A). Then, to analyze the relevance of IL-17 signaling on human tumor progression, we selected two cancer types Skin Cutaneous Carcinoma (SKCM) and Acute Myeloid Leukemia (LAML) that presented high and low IL-17RC transcript levels, respectively, and shared tissue origins with our cancer experimental models. We compared the expression of *Il17ra, Il17rc* and *Il17rd* in patient samples from both tumor types and observed that the three transcripts were detectable in samples from both tumor types, though samples from SKCM showed lower levels of *Il17ra* transcripts and higher levels of *Il17rc* and *Il17rd* transcripts than LAML samples (Figure 8A). We next investigated possible correlation between the level of IL-17R subunit expression and patient survival. We observed that within the SKCM cohort, patients with high expression of IL-17RA showed longer survival than patients with low IL-17RA expression (Figure 8B). This difference in the overall survival related to the level of IL-17RA expression was not observed in the LAML cohort (Figure 8C). We also noted that the expression levels of *Il17rc* or *Il17rd* in SKCM and LAML were not associated with differences in survival (Supplementary Figure 6B). Altogether, this analysis suggests that IL-17RA signaling is protective in tumors like SKMC that express IL-17RA together with high levels of IL-17RC (and IL-17RD), in accordance with the data obtained from our experimental cancer models.

**Figure 8.**
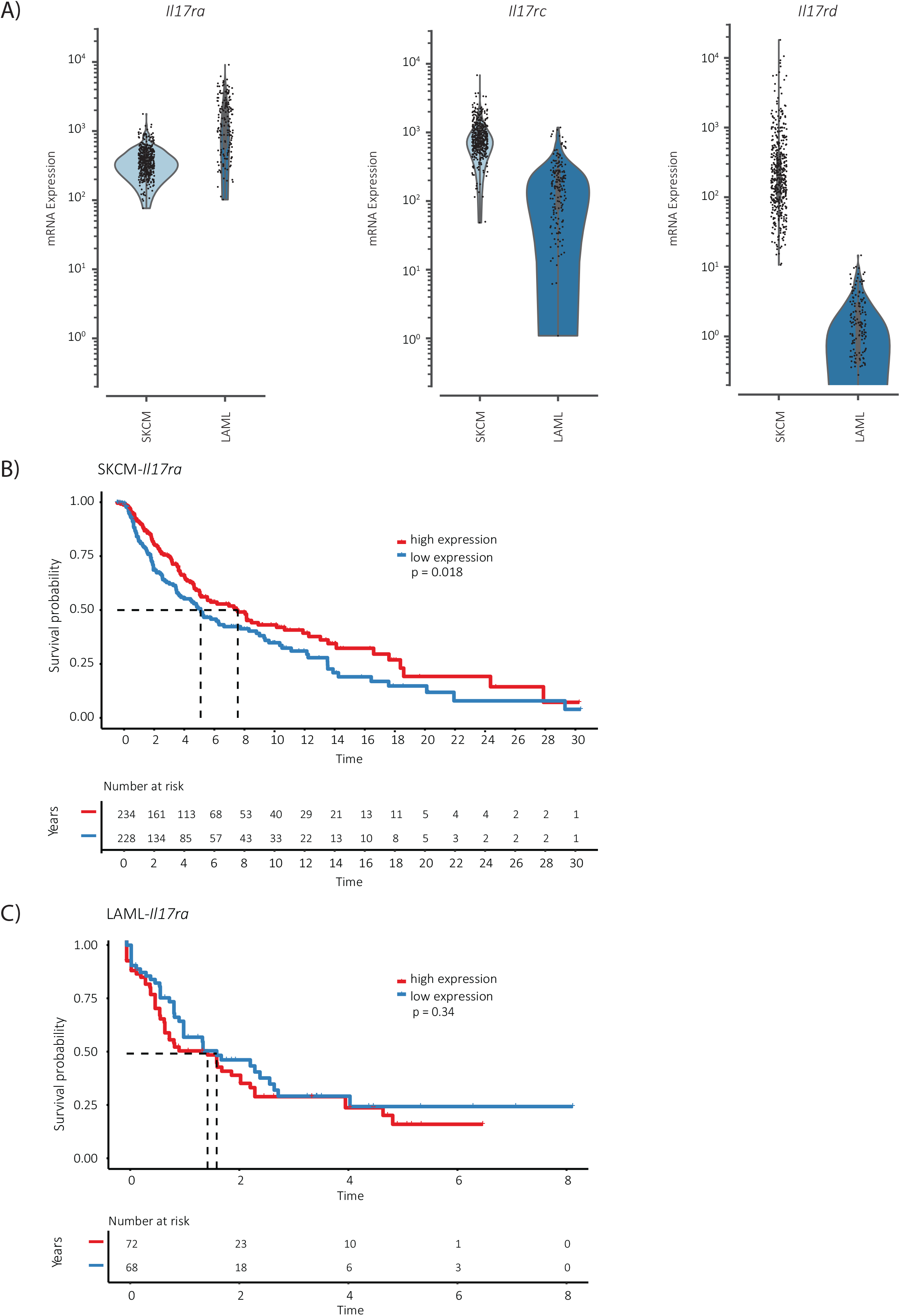
High IL-17RA expression correlates with increased survival probability in patients with melanoma (SKCM) but not in patients with leukemia (LAML). **(A)** Level of *il17ra, il17rc* and *il17rd* mRNA amount in tumor samples from SKCM and LAML patients. **(B-C)** Kaplan Meier curves of survival probability in patient cohorts with SKCM **(B)** and LAML **(C)** tumors that showed high (red) and low (blue) expression of the IL-17RA subunit. Data of gene expression and overall survival (defined as time to death) was obtained from the Cancer Genome Atlas portal. The statistical significance of the overall survival curves stratified according to high and low expression of IL-17RA for each tumor type was determined by a Log-Rank test (R package “survival”).

## DISCUSSION

IL-17 mediated responses have been linked to dramatically different net outcomes in cancer. Therefore, the scientific community is still debating the exact roles of this cytokine in tumor progression and its potential as a novel therapeutic target to treat malignancies (5, 12). In this work, we established that tumors of mesenchymal/epithelial origin (i.e melanoma and fibrosarcoma) exhibited an increased tumor progression while hematopoietic tumors (i.e. lymphoma and leukemia) showed reduced growth in hosts with deficient IL-17 signaling. Lack of IL-17RA or IL-17A/F expression in the hosts had several tumor-specific effects influencing the production of inflammatory, angiogenic and metastatic mediators and the immune cell infiltrate. In contrast, deficiencies in host IL-17 signaling molecules limited tumor-specific CD8+ T cell immunity independently of the cancer model and the final impact on tumor growth. Tumor-specific effects correlated with distinct patterns of IL-17R subunit expression in the tumor cells that led to particular transcriptional responses to IL-17. Specifically, the expression of IL-17RC emerged as a pivotal determinant in the canonical signaling downstream IL-17 stimulation that likely orchestrates global anti-tumor functions. Accordingly, high IL-17RA expression was associated with increased survival in cohorts of patients with tumors such as melanoma, characterized by a high IL-17RC expression.

In agreement with our results, several studies have associated IL-17-mediated responses with an improved control of this murine melanoma model (24, 52, 53). In contrast, others reported that IL-17 globally promotes the growth of B16 tumors (54–56). We hypothesize that tumor immunogenicity may be one pivotal aspect to reconcile the contrasting observations regarding IL-17 roles in cancer. In implanted tumor models, immunogenicity may increase due to the expression of foreign antigens such as OVA or SIY, and influence the robustness of antitumor responses, tipping the balance of IL-17 function towards tumor elimination. Evidences supporting this notion come from reports indicating that IL-17 can stimulate the anti-cancer response and tumor elimination in immunocompetent mice ((24, 27, 57), but, on the contrary, this cytokine promotes angiogenesis and tumor growth in immunocompromised mice where the adaptive immune response is missing (58). Notably, studies where a pro-tumor role was reported for IL-17 used parental B16 cells (54–56), which are known to generate tumors with a relatively poor T cell infiltration that may underline a reduced immunogenicity (59, 60). In this regard, tumors generated by parental B16 cells result more aggressive and faster growth rate in comparison to tumors generated by the B16.SIY cell line variant used in our work (31, 61). Noteworthy, we determined that IL-17 signaling participates in the control of not only the B16.SIY melanoma but also the MC57.SIY fibrosarcoma. Although no studies have reported IL-17 roles on fibrosarcoma progression so far, it is known that MC57.SIY rejection depends on tumor antigen cross-presentation and CD8+ T cell recognition (30, 62). In overall, these data suggest that the presence of immune cells able to recognize tumor antigens within the tumor microenvironment may be a fundamental requirement for IL-17 anti-tumor functions. In clear contrast, we found that the IL-17A/F:RA pathway promoted the growth of hematopoietic tumors generated by the EL4.SIY and C1498.SIY cell lines. Although no previous studies evaluated the role of IL-17 in the C1498 acute myeloid leukemia model, our findings are coincident with previous reports describing positive effects of this cytokine in the development of the EL4 lymphoma. In this tumor context, IL-17 signaling was shown to favor immunosuppressive tumor microenvironments (18, 63) and promote angiogenesis through stromal cell activation (63). Thus, the dominant IL-17 effector mechanisms at place in these settings may be related to the exacerbation of pro-angiogenic and pro-tumorigenic pathways by direct signaling on cells within the tumor microenvironment.

In agreement with the role of IL-17 in the modulation of molecules such as VEGF, CXCL1, CXCL8, CXCL9 and CXCL10 among others (18, 23, 58), we determined that host IL-17RA or IL-17A/F deficiency affected the expression of several inflammatory and angiogenic proteins within the melanoma and lymphoma niches. We identified not only proteins such as MMP-3 and MMP-9 that are known IL-17 targets (45) as well as PTX3 and Serpin F1 that were shown to be modulated by IL-17 in other cell contexts (64, 65), but also other mediators like IGFBP2 which has not been previously linked to this cytokine pathway. Although exploratory in nature, our studies highlight that IL-17 signaling has, either directly or indirectly, a profound tumor-specific impact on the tumor microenvironment. Further studies are needed to establish a link between the modulation of particular angiogenic and inflammatory proteins and the ultimate outcome of deficient IL-17 signaling in the hosts. Together with the modulation of pro- and anti-tumor mediators, IL-17 may affect tumor progression by influencing the infiltration of immune cells. It has been reported that IL-17A restrains tumor growth by promoting a cytotoxic microenvironment through the development and recruitment of anti-tumor myeloid cells, antigen-presenting cells and activated effector T cells (5, 23). In addition, the IL-17 pathway sustains NK cell and CD8+ T cell responses in the context of infections (47, 66, 67), mechanisms likely operating also in cancer. On the contrary, IL-17 may lead to tumor permissive microenvironments by recruiting a myeloid compartment with immunosuppressive functions (18, 19, 68) and inducing regulatory molecules like PD-L1 (51). Surprisingly, we established that lack of IL-17RA or IL-17A/F expression in murine hosts had a minor or no impact in monocyte and neutrophil tumor infiltration, although it led to alterations in lymphoid subsets. Thus, tumor-specific and effector CD8+ T cells were decreased in B16.SIY tumor cell infiltrates from both IL-17 deficient mice while exhausted-like, PD1 and LAG3 co-expressing CD8+ T cells were not affected. These results contrast with those showing that, in the absence of CD4+ T cell help, Tc17 cells favored the generation of terminally exhausted CD8+ T cells and therefore IL-17 was linked to a poorer tumor growth control in a murine model of metastatic melanoma (48). Depletion of CD4+ T cells may emerge as a source of divergence as it is a widely reported approach to induce high percentage of exhausted CD8+ T cells in models of chronic LCMV infections (69–71) and to improve anti-tumor response in certain cancer models likely by targeting regulatory T cells (72, 73). Therefore, in settings where CD4+ T cell depletion unlock intra-tumor immunoregulation, IL-17 signaling may predominantly contribute to chronic inflammation and this, in turn, potentiates CD8+ T cell exhaustion and becomes detrimental in the global context of the tumor. Notably, reduced tumor-specific CD8+ T cells were also detected in EL4.SIY tumors developed in IL-17RA KO hosts but not in IL-17A/F DKO mice in which IL-17A and IL-17F production by the lymphoma itself could compensate, at least in part, for the lack of endogenous IL-17A/F production, reverting mouse phenotype. Overall, our data highlight that the positive effect of IL-17 on CD8+ T cell immunity may be a general mechanism operating not only in infections as we reported previously (47, 67), but also in cancer independently of the tumor type and not necessarily linked to better tumor progression. Finally, we determined that IL-17A/F:RA pathway deficiencies in the host also affected tumor B cell infiltration in opposite directions depending on the tumor type. Although IL-17 was previously shown to influence the development and recruitment of B cells to the inflammatory site in an esophageal cancer model (74), the relevance of this effect remains unexplored. Altogether, our findings may reorientate the evaluation of the immune cell populations acting downstream the IL-17 pathway in different cancer contexts, with a focus in lymphoid subsets.

Considering the preponderance of tumor cells, it is likely that their differential response to IL-17 signaling may be a critical factor shaping a distinct microenvironment that becomes more permissive or restrictive for tumor growth. In this regard, we determined that non-hematopoietic tumor cells (i.e. B16.SIY and MC57.SIY) expressed IL-17RA, IL-17RC and IL-17RD and do not produce IL-17A or IL-17F, while in contrast, the hematopoietic cell lines (i.e. EL4.SIY and C1498.SIY) express only IL-17RA and IL-17RD and produce both cytokines. These results are coincident with the available literature which indicates that the expression of IL-17RA and IL-17RD is ubiquitous in physiological contexts, whereas IL-17RC is expressed to a major extent in non-hematopoietic epithelial and mesenchymal cells (75–78), and that IL-17A and IL-17F production is associated with immune cells (79). Noteworthy, differences in the expression profiles of the IL-17R subunits among the tumor cell lines showed biological relevance by imprinting different response upon IL-17 stimulation. In this sense, only epithelial/mesenchymal cells that expressed the three IL-17R subunits, but not tumor cells of hematopoietic origin that lacked IL-17RC, showed a conventional response to IL-17A or IL-17F stimulation with a moderate phosphorylation and nuclear translocation of NF-kB and an IL-17-associated transcriptional signature (80). Remarkably, we determined that IL-17 activated gene transcription also in the hematopoietic EL4.SIY cells in a response characterized by the induction of biological cellular processes not typically associated with IL-17 functions. Some of the processes like “Factors promoting carcinogenesis in vertebrates” are directly associated with cancer while others such as the “Axonal and synaptogenic signaling” have also been associated with tumor promotion given their role in apoptosis inhibition, cell migration and neural network formation that sustain tumor growth (81–83). So far, studies aimed at evaluating the role of IL-17 in cancer have focused on the expression of IL-17RA or IL-17RC separately (84, 85) without a systematic analysis of the heterodimer relevance, with few exceptions (63). In addition, under the idea that IL-17 can only signal through the canonical IL-17RA/RC receptor complex, the evaluation of IL-17RD expression in tumors in the context of IL-17 signaling has not been incorporated yet (86). Within this incomplete scenario, our results support the notion that different types of tumors respond to IL-17 by activating distinct signaling pathways and signatures according to their profiles of IL-17R subunit expression and this may underlie the reported dual effects of this cytokine in cancer progression. Remarkably, IL-17RC blockade markedly changed the response to IL-17 stimulation in B16.SIY cells, imprinting an immunoregulatory and likely pro-tumor profile similar to that observed in EL4.SIY cells where IL-17 signals through the heterodimer formed by IL-17RA and IL-17RD (or other related receptor family) due to the lack of IL-17RC expression. Whether these changes detected *in vitro* have the predicted effect *in vivo* needs further evaluation using blockade approaches or IL-17RC deficient cells lines. In this regard, Yan et al. (87) showed that IL-17RC silencing alters *in vitro* basal proliferation and *in vivo* tumor growth in a tumor-type dependent manner. Remarkably, the distinctive functions of IL-17RC signaling were not attributed to the IL-17RC expression levels but rather to differences in the basal expression of downstream molecules involved in intracellular signaling and tumor cell proliferation. Interestingly, another layer of regulation of the IL-17 pathways is provided by the different signaling capacity of the cytokines of the family. In this regard, IL-17F can signal through an IL-17RC/RC homodimer without the requirement of IL-17RA in humans (10). Furthermore, the signaling of IL-17A but not of IL-17F through the IL-17RA/IL-17RD pathway is involved in the pathogenesis of psoriasis (9).

Differences in mice and humans regarding expression and binding affinities of cytokines and receptors from the IL-17 pathway (77), may limit the extrapolation of experimental results to clinical settings. In general terms, IL-17RA and IL-17RD are expressed at variable levels in all human tissues under physiological conditions (75–78). In contrast, IL-17RC is expressed to a greater extent in the liver, prostate, skin, colon, stomach, lung and thyroid, while in lymph nodes, bone marrow and bladder its expression is lower (77). Accordingly, our analysis of TCGA-based data demonstrated significant heterogeneity in the expression of IL-17RA, IL-17RC and IL-17RD transcripts across cancer types. In this context, we attempted to correlate the subunit expression profiles with the effect of the IL-17 pathway on human tumor progression and selected the melanoma (SKCM) and acute myeloid leukemia (LAML) tumors. Differences in tumor IL-17RA expression levels correlated with differences in overall survival time in SKCM patients with high tumor IL-17RC expression but not in LAML patients with low tumor IL-17RC expression while the expression levels of IL-17RC itself or IL-17RD were not associated with differences in survival in any of the cohorts. Recently, deletion of IL17RA or IL-17RC in the colon was associated with advances stages and a worse prognosis in colorectal cancer prognosis (88). In addition, other studies evaluating the role of the IL-17 pathway in cancer patients highlighted that IL-17A levels were more frequently associated with a worse prognosis although an association with improved survival was also reported not only for tumors of different histological origin but also for the same tumor type (11). Once again, these evidences underline the critical need to study each cancer settings in order to position the IL-17 pathway as a putative target of personalized antitumor therapies.

In conclusion, our results show that the IL-17A/IL-17R pathway has diverse concomitant effects on different components of the tumor microenvironment, influencing tumor development. It addition, IL-17 signaling is required to generate robust specific effector CD8+ T cell responses that would participate in the control of tumor growth. At the same time, IL-17A triggers various responses in different tumor cells dictating inflammatory processes that orchestrate the immune infiltrate but also activating cellular events that enhance tumor growth. These differential downstream pathways correlate with particular expression of IL-17R subunits with IL-17RC emerging as a critical molecular determinant of the global pro- or anti-tumor effect of IL-17 signaling. In this sense, our work proposes the study of the expression profile of different IL-17R subunits together with this pathway’s cytokines to rationally define the usefulness of therapies aimed at controlling the IL-17 axis in personalized antitumor immunotherapies.

## Acknowledgements

We thank the staff from Cytometry Core, Animal, Cell culture and Molecular biology facilities from CIBICI-CONICET at Facultad de Ciencias Químicas, UNC. We also like to acknowledge Tomas Gajewski, Mercedes Fuertes and Ximena Raffo who provided the cell lines and protocols for cell culture. CR, SB, and SNB thank CONICET for the fellowship awarded. LF, DGC, CM, AG and EVAR are members of the Scientific career of CONICET. EP and JT are supported by the LabEx DCBIOL (ANR-10-IDEX-0001-02 PSL; ANR-11-LABX-0043); SIRIC INCa-DGOS-Inserm 1255; and Center of Clinical Investigation (CIC IGR-Curie 1428).

## Supplementary Material

### Supplementary Methods

#### Antibodies used for flow cytometry

**Supplementary Table 1:** Tumor infiltrating cells subpopulations Panel.

| Antigens | Fluorochrome | Source | Identification number | Dilution |
| --- | --- | --- | --- | --- |
| B220 | Pacific Blue | Biolegend | Cat: 103227 | 1/200 |
| MHCII | Super Bright 600 | Thermofisher | Cat: 63-5321-82 | 1/500 |
| CD11B | SUPER BRIGHT 645 | Thermofisher | Cat: 64-0112-80 | 1/300 |
| Ly-6G/Ly-6C | Super Bright 702 | Thermofisher | Cat: 67-5931-82 | 1/200 |
| Ly-6C | APC | eBioscience | Cat: 17-5932-82 | 1/300 |
| CD45 | AF700 | Thermofisher | Cat: 56-0451-82 | 1/100 |
| F4/80 | APC-Cy7 | Biolegend | Cat: 123118 | 1/200 |
| CD49b | PE | eBioscience | Cat: 12-5971-83 | 1/200 |
| CD4 | PE-CF594 | BD | Cat: 562285 | 1/300 |
| CD19 | PE-Cy5 | eBioscience | Cat: 15-0193-82 | 1/300 |
| CD11c | PE-Cy7 | eBioscience | Cat: 25-0114-82 | 1/100 |
| CD8 | PE-Cy5.5 | eBioscience | Cat: 35-0081-82 | 1/300 |
| CD3 | Brillant Violet 785 | Biolegend | Cat: 100355 | 1/100 |

**Supplementary Table 2:** CD8 phenotype Panel.

| Antigens | Fluorochrome | Source | Identification number | Dilution |
| --- | --- | --- | --- | --- |
| LAG3 | PerCP-eF710 | eBioscience | Cat: 46-2231-82 | 1/100 |
| CD127 | Brillant Violet 421 | BD | Cat: 566377 | 1/100 |
| CXCR3 | Brillant Violet 510 | Biolegend | cat: 126528 | 1/100 |
| CD62L | Super Bright 600 | eBioscience | Cat: 63-0621-82 | 1/200 |
| CD4 | Brillant Violet 650 | BD | Cat: 563232 | 1/300 |
| B220 | Brillant Violet 711 | Biolegend | Cat: 103255 | 1/200 |
| CD3 | Brillant Violet 785 | Biolegend | Cat: 100355 | 1/100 |
| CD69 | APC | BD | Cat: 560689 | 1/100 |
| CD45 | APC-Cy7 | BD | Cat: 557659 | 1/400 |
| H-2 Kb (SIYRYYGL) | PE | Immudex | Cat: JD2164 | 1/10 |
| KLRG1 | PE-eFluor 610 | eBioscience | Cat: 61-5893-82 | 1/300 |
| CD44 | PE-Cy5 | BD | Cat: 553135 | 1/400 |
| CD8 | PE-C5.5 | eBioscience | Cat: 35-0081-82 | 1/300 |
| PD1 | PE-Cy7 | eBioscience | Cat: 25-9985-82 | 1/100 |

**Supplementary Table 3:** Other antibodies and Reagents.

| Reagent | Source | Identification | Dilution |
| --- | --- | --- | --- |
| Fixable Viability Stain 700 | BD | Cat: 564997 | 1/500 |
| Fixable Aqua Dead Cell Stain Kit, for 405 nm excitation | Invitrogen | Cat: L34957 | 1/500 |
| IL-17RA - PE | eBiosciences | 12-7182-80 | 1/10 |
| IL-17RC | R&D | BAF2270 | Protocol |
| IL-17RD | R&D | BAF2276 | Protocol |
| Streptavidin - APC | Biolegend | 405207 | 1/300 |

#### Antibodies used for western blot

**Supplementary Table 4:** Western Blot antibodies.

| Antigen | Company | Catalog | Dilution |
| --- | --- | --- | --- |
| IL-17RD-biotin | R&D | BAF2276 | 1/2000 |
| Phospho-p44/42 (ERK1/2) | Cell signaling | #4370 | 1/500 |
| p44/24 (ERK1/2) | Cell signaling | #9102 | 1/500 |
| Phospho-NF-κB p65 | Cell signaling | #3033 | 1/500 |
| NF-κB p65 | Cell signaling | #8242 | 1/500 |
| Phospho-p38 | Santa Cruz | #sc-7975 | 1/500 |
| GAPDH | Santa Cruz | #sc-25778 | 1/1000 |
| PARP-1 | Santa Cruz | #sc-7150 | 1/200 |

#### Taqman probes used for qRT-PCR

**Supplementary Table 5:**
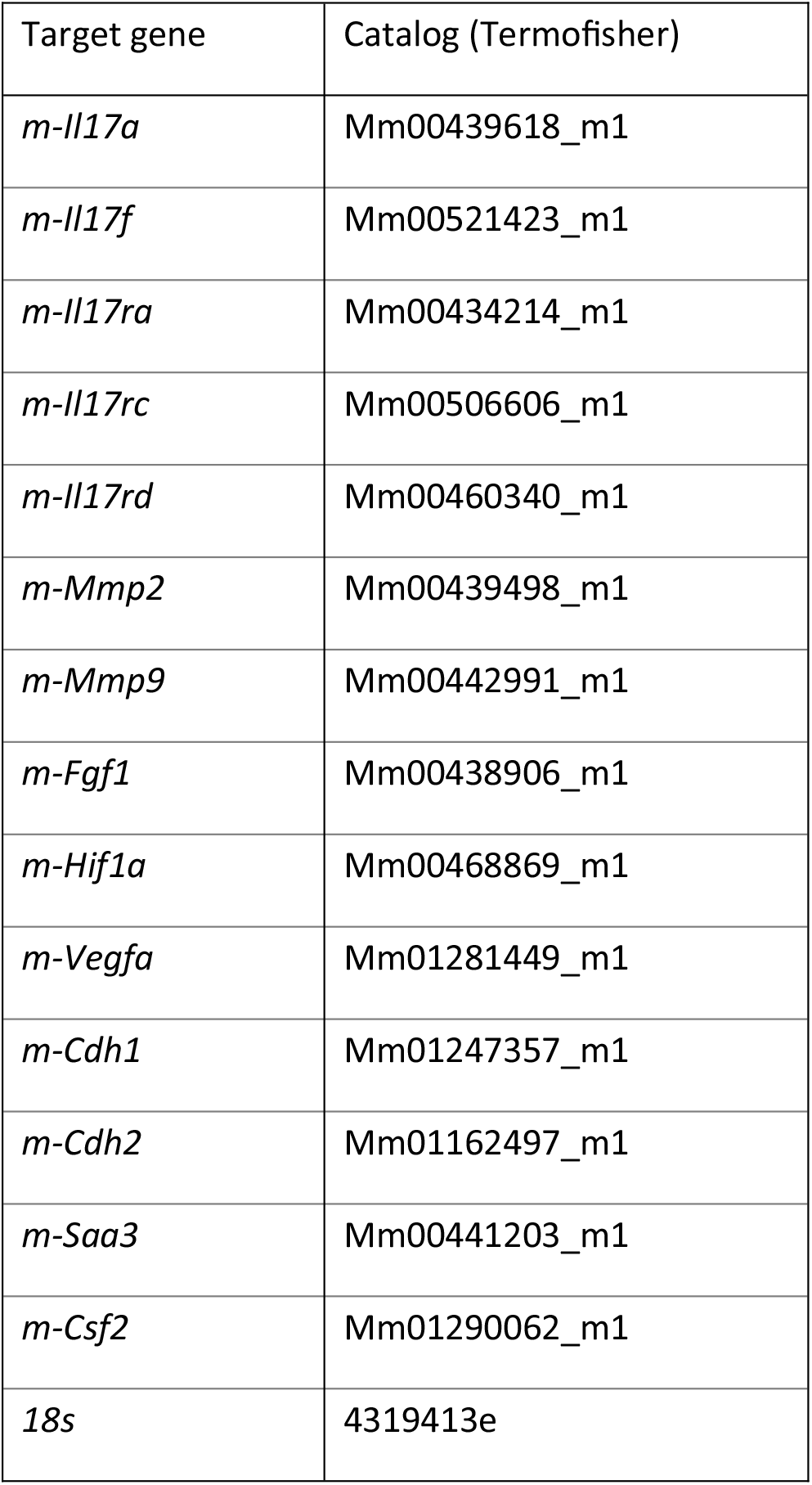
Taqman probes.

### Supplementary figures

**Supplementary Figure 1.**
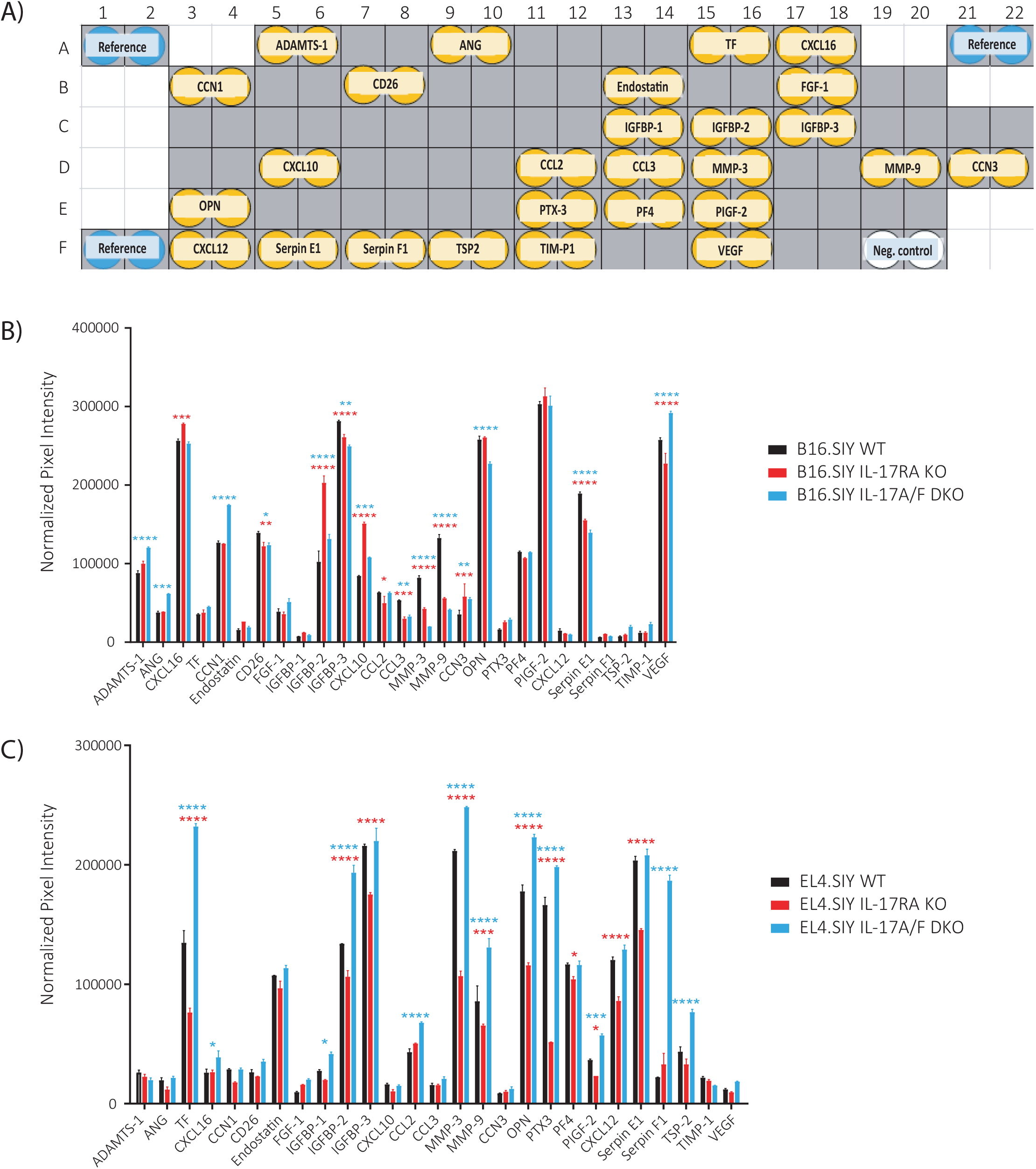
Analysis of the expression of inflammation- and angiogenesis-related proteins using the Proteome Profiler Mouse Angiogenesis Array Kit (ARY015, R&D). We assessed the relative levels of inflammation- and angiogenesis-related proteins in the B16.SIY and EL4.SIY tumors lysates collected at 17-18dpi from WT, IL-17RA KO and IL-17A/F DKO mice. **(A)** Representation of the quantification mask of the 27 proteins detected and the reference sport. FIJI sotfware was used to build the mask and to assess the pixel intensity of the spots (represented in yellow) **(B-C)** Statistical analysis of the pixel intensity of the proteins detected normalized to the reference spots, determined in lysates of B16.SIY **(B)** and EL4.SIY **(C)** tumors from IL-17RA KO (red bars), IL-17A/F DKO (blue bars) and WT (black bars) mice. p-values were calculated by two-way ANOVA and subsequent Dunnett’s multiple comparison. * p < 0.05; ** p < 0.01; *** p < 0.001; **** p < 0.0001.

**Supplementary Figure 2.**
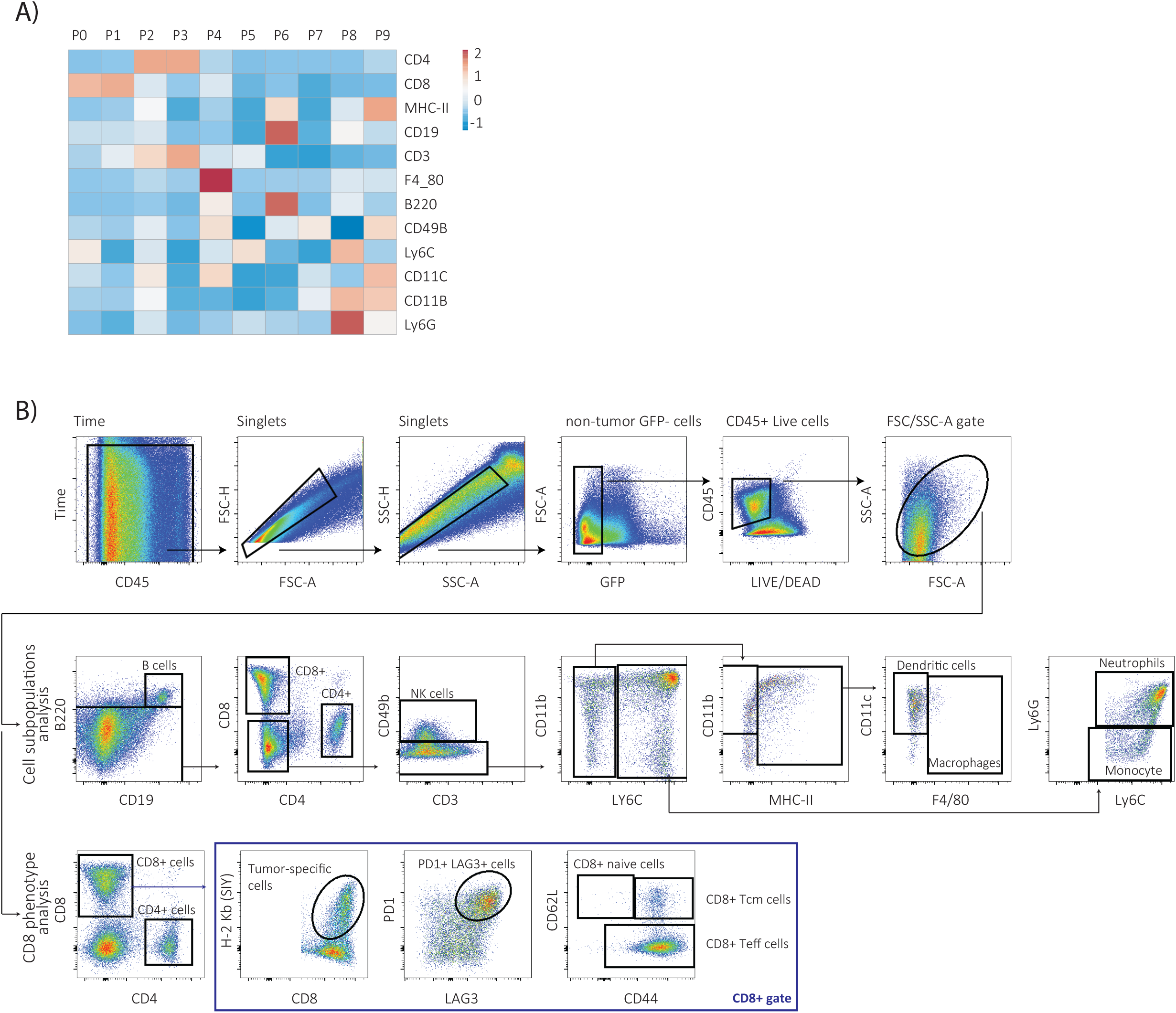
Strategy of multiparametric flow cytometry analysis of tumor-infiltrating immune cell subsets. B16.SIY and EL4.SIY tumors developed in WT, IL-17RA KO and IL-17A/F DKO mice were collected at endpoint, disaggregated using DNAsa and Colagenase IV for 30 min, washed and then stained to identify distinct tumor-infiltrating immune cells. **(A)** The heat-map shows the relative mean fluorescence intensity of the 12 markers used to identify different cell clusters within the CD45+ gate in B16.SIY and EL4-SIY tumors obtained from WT mice using unsupervised analysis combining UMAP followed by FlowSOM. **(B)** Gating strategy for the supervised analysis of B16.SIY and EL4.SIY tumorinfiltrating immune and myeloid subpopulations and tumor-infiltrating CD8+ T cells phenotype.

**Supplementary Figure 3.**
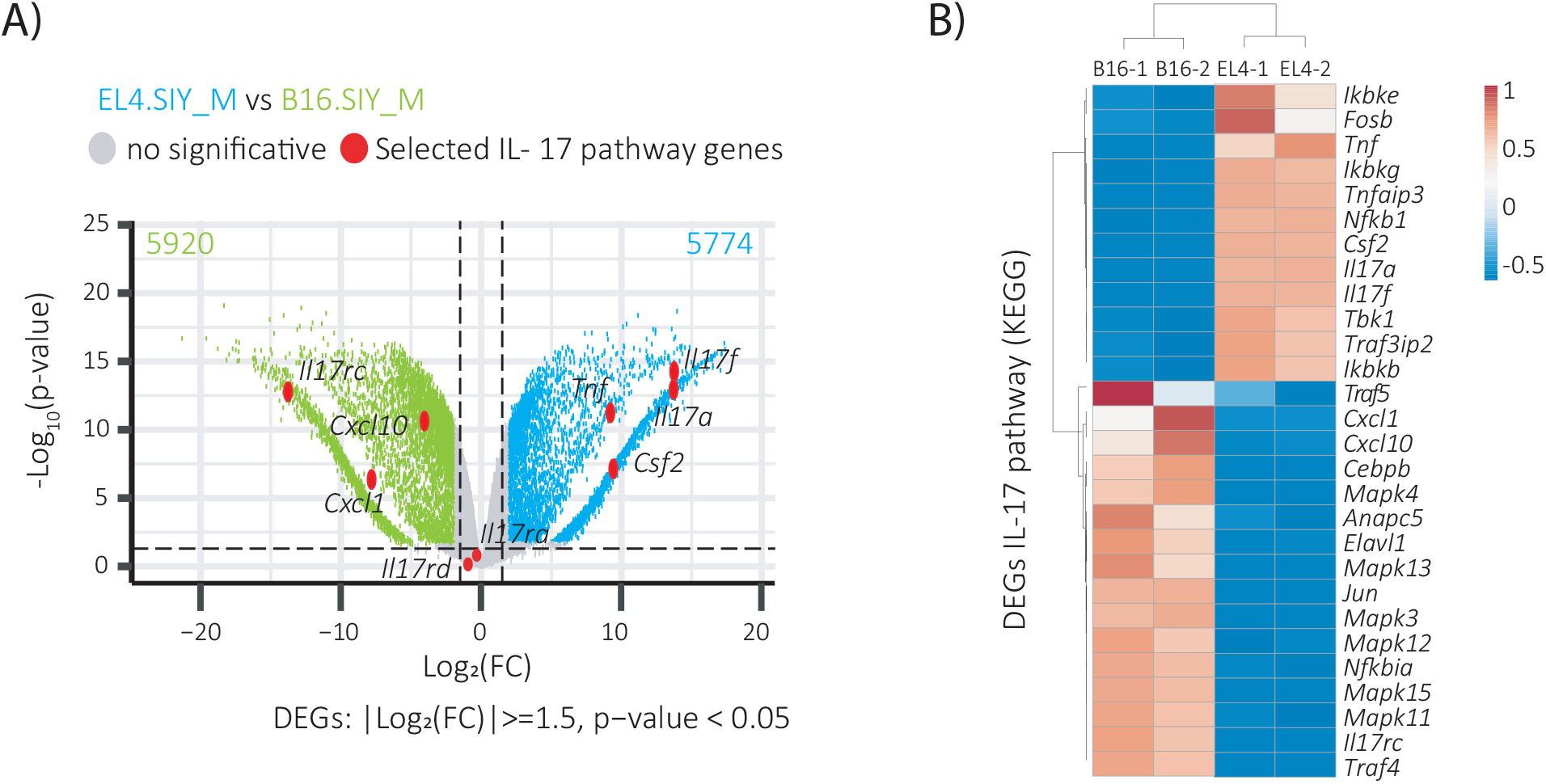
B16.SIY and EL4.SIY cells show differences in the basal expression of genes associated with the IL-17 pathway. **(A)** The volcano plot depicts significant differentially expressed genes (DEGs) between B16.SIY (green) and EL4.SIY (blue) cells. Genes without significant changes are shown in grey. Selected genes belonging to the IL-17 pathway are highlighted in red. **(B)** Heat map showing the complete list of DEGs in EL4.SIY versus B16.SIY cell lines belonging to the IL-17 pathway in the KEGG database. n=2 replicates per group. nd: non detectable. One-way Anova was used for statistical analysis. * p value <0,05 ** p value<0,01 ***p value<0,001 ****p value < 0,0001.

**Supplementary Figure 4.**
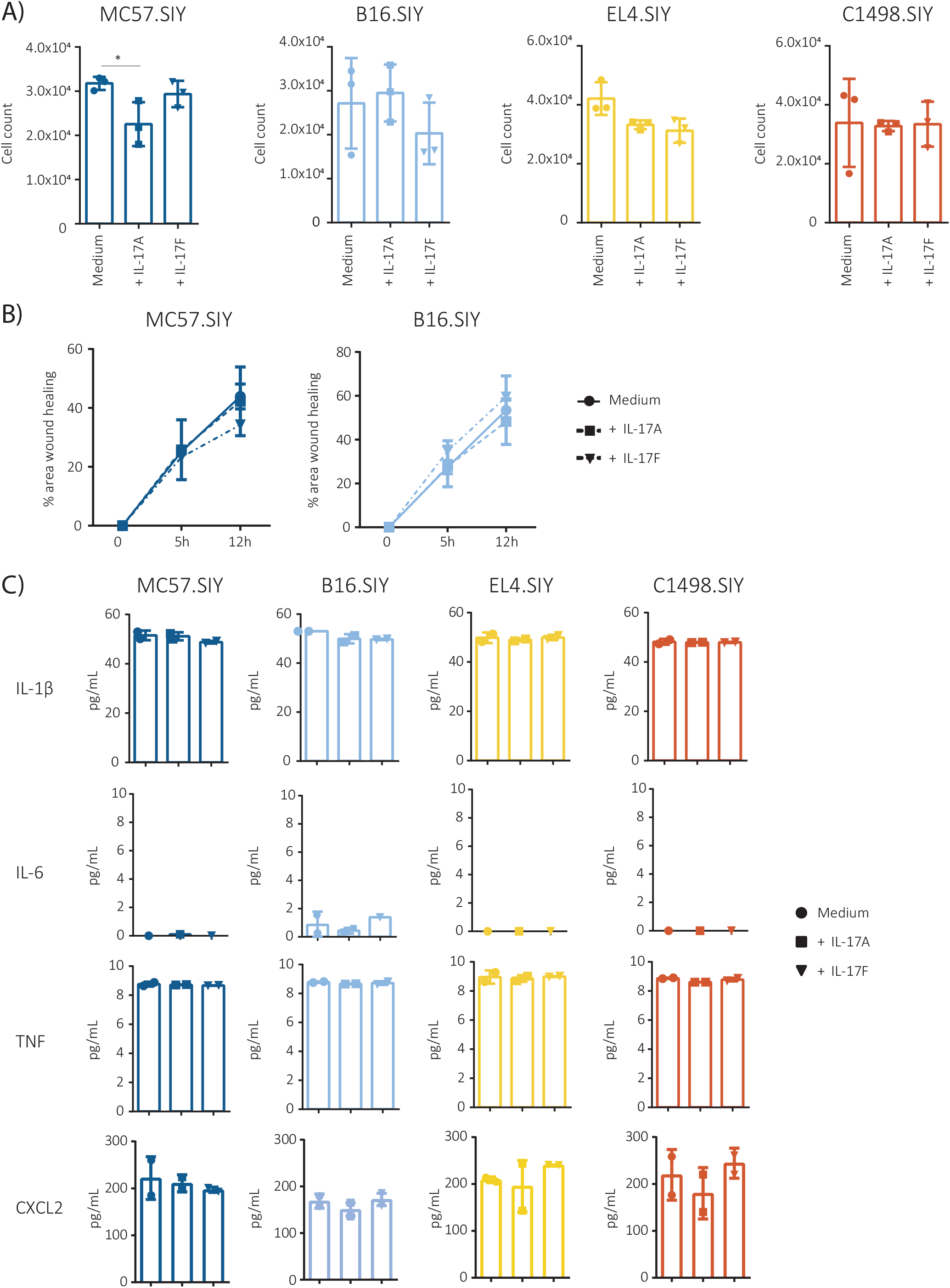
In vitro stimulation with IL-17A or IL-17F had no significant effect on the proliferation, migration and production of inflammatory cytokines by the tumor cell lines evaluated. MC57.SIY, B16.SIY, EL4.SIY and C1498.SIY cells were incubated in serum reduced conditions and stimulated with IL-17A or IL-17F (200 ng/ml) during the indicated times. **(A)** Cell numbers determined after 24 h of culture in the indicated conditions. **(B)** Percentage of wound closure at different times after culture of adherent B16.SIY and MC57.SIY cells in the indicated conditions. **(C)** Concentration of IL-1β, IL-6, TNF and CXCL2 determined by ELISA in the supernatants of the different tumor cell lines cultured during 48h in the indicated condition. Results are shown as the mean ± SD (N=2-3) for each condition. p-values were calculated by unpaired two-tailed Student’s t-test. * p < 0.05; ** p < 0.01; *** p < 0.001; **** p < 0.0001.

**Supplementary Figure 5.**
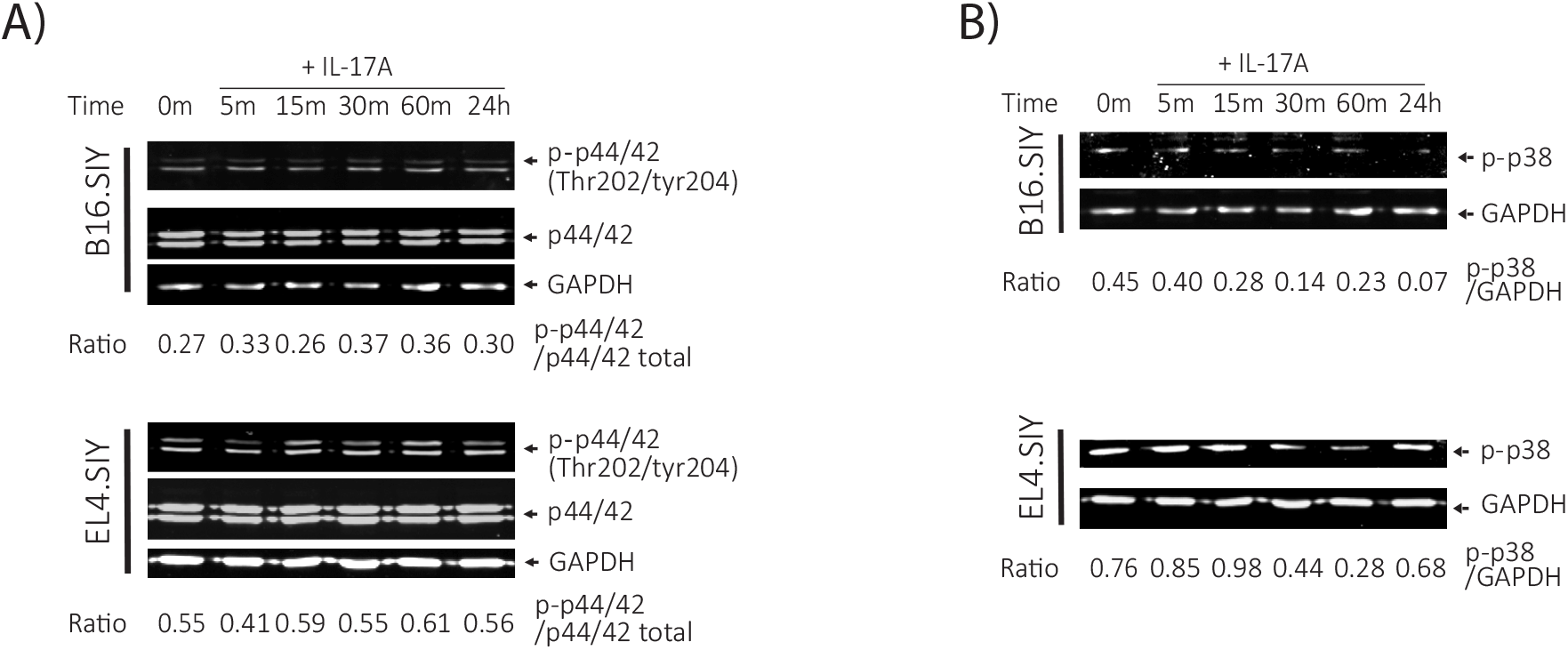
In vitro stimulation of B16.SIY and EL4.SIY tumor cells with IL-17 did not result in ERK and JNK phosporilation. **(A-B)** Phosphorylation of Erk (p44/42) **(A)** and p38 **(B)** MAPKs of B16.SIY and EL4.SIY cells after stimulation with IL-17A (200 ng/mL) for 5, 15, 30, 60 minutes (min) and 24 h determined by western blot in whole cell lysates. GAPDH protein expression was used as loading control. The indicated ratio values between the relative expression of the indicated proteins were determined using the FIJI software.

**Supplementary Figure 6.**
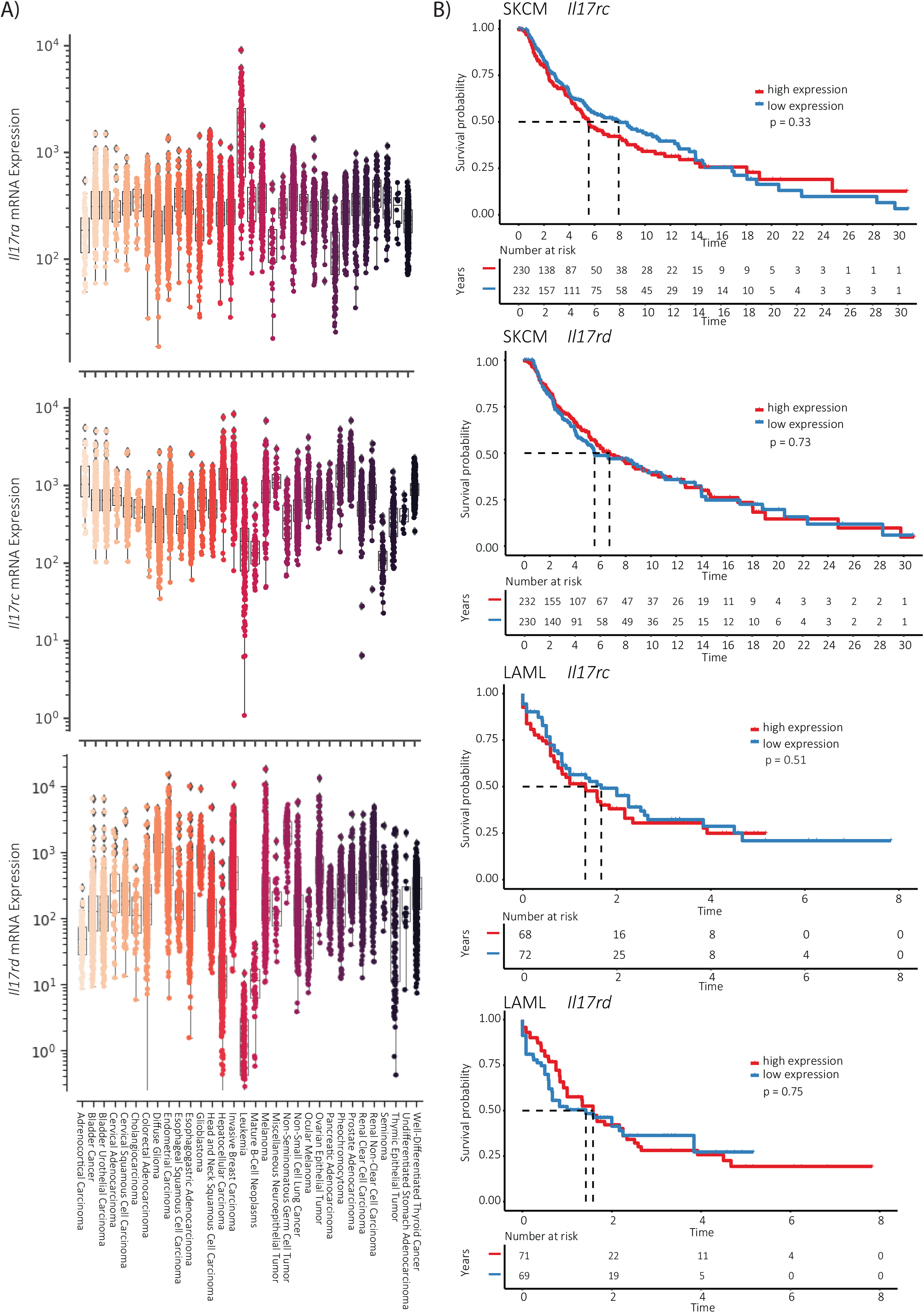
Il17rc or Il17rd expression levels in SKCM and LAML were not associated with differences in overall survival time in the patient cohorts. **(A)** Analysis of *Il17ra, Il17rc*, and *Il17rd* mRNA expression from various tumors selected from the TGCA project. Data were downloaded from the c-BioPortal platform. Quantification values from mRNA expression were obtained with the RSEM tool and were batch-normalized. **(B)** Kaplan Meier curves of survival probability in patient cohorts with SKCM and LAML tumors with high (red) and low (blue) *Il17rc* and *Il17rd* subunit expression. The statistical significance of the overall survival curves (defined as time to death), stratified by the aforementioned groups for each tumor type, was determined by a Log-rank test (R survival package).

